# *V. longisporum* elicits media-dependent secretome responses with a further capacity to distinguish between plant-related environments

**DOI:** 10.1101/2020.02.11.943803

**Authors:** Miriam Leonard, Anika Kühn, Rebekka Harting, Isabel Maurus, Alexandra Nagel, Jessica Starke, Harald Kusch, Oliver Valerius, Kirstin Feussner, Ivo Feussner, Alexander Kaever, Manuel Landesfeind, Burkhard Morgenstern, Dörte Becher, Michael Hecker, Susanna A. Braus-Stromeyer, James W. Kronstad, Gerhard H. Braus

**Affiliations:** Department of Molecular Microbiology and Genetics, Institute of Microbiology and Genetics and Goettingen Center for Molecular Biosciences (GZMB), University of Goettingen, Goettingen, Germany.; Department for Plant Biochemistry, Albrecht-von-Haller-Institute for Plant Sciences and Goettingen Center for Molecular Biosciences (GZMB), University of Goettingen, Goettingen, Germany.; Department of Bioinformatics, Institute of Microbiology and Genetics and Goettingen Center for Molecular Biosciences (GZMB), University of Goettingen, Goettingen, Germany.; Department of Microbial Physiology, Institute for Microbiology, Ernst-Moritz-Arndt-Universität Greifswald, Greifswald, Germany.; Michael Smith Laboratories, Department of Microbiology and Immunology, University of British Columbia, Vancouver, Canada.

## Abstract

Verticillia cause a vascular wilt disease affecting a broad range of economically valuable crops. The fungus enters its host plants through the roots and colonizes the vascular system. It requires extracellular proteins for a successful plant colonization. The exoproteome of the allodiploid *Verticillium longisporum* was analyzed upon cultivation in different media. Secreted fungal proteins were identified by label free LC-MS/MS screening. *V. longisporum* induced two main secretion patterns. One response pattern was elicited in various non-plant related environments. The second pattern includes the exoprotein responses to the plant-related media, pectin-rich simulated xylem medium and pure xylem sap, which exhibited similar but additional distinct features. These exoproteomes include a shared core set of 223 secreted and similarly enriched fungal proteins. The pectin-rich medium significantly induced the secretion of 144 proteins including a number of pectin degrading enzymes, whereas xylem sap triggered a smaller but unique fungal exoproteome pattern with 32 enriched proteins. The latter pattern included proteins with domains of known effectors, metallopeptidases and carbohydrate-active enzymes. The most abundant and uniquely enriched proteins of these different groups are the necrosis and ethylene inducing-like proteins Nlp2 and Nlp3, the cerato-platanin proteins Cp1 and Cp2, the metallopeptidases Mep1 and Mep2 and the CAZys Gla1, Amy1 and Cbd1. Deletion of the majority of the corresponding genes caused no phenotypic changes during *ex planta* growth or invasion and colonization of tomato plants. However, we discovered that the *NLP2* and *NLP3* deletion strains were compromised in plant infections. Overall, our exoproteome approach revealed that the fungus induces specific secretion responses in different environments. The fungus has a general response to non-plant related media whereas it is able to fine-tune its exoproteome in the presence of plant material. Importantly, the xylem sap-specific exoproteome pinpointed Nlp2 and Nlp3 as single effectors required for successful *V. dahliae* colonization.

**Author Summary:** *Verticillium* spp. infect hundreds of different plants world-wide leading to enormous economic losses. Verticillium wilt is a disease of the vasculature. The fungus colonizes the xylem of its host plant where it exploits the vascular system to colonize the whole plant. Therefore, the fungus spends part of its lifetime in this nutrient-low and imbalanced environment where it is inaccessible for disease control treatments. This lifestyle as well requires the fungus to react to plant defense responses by secreting specific effector molecules to establish a successful infection. We addressed the differences in media-dependent secretion responses of *Verticillium longisporum*. We identified a broad response pattern induced by several media, and a similar response (but with some distinct differences) for the plant-related environments: the pectin-rich medium SXM and xylem sap from the host rapeseed. Importantly, we show that the necrosis and ethylene inducing-like proteins Nlp2 and Nlp3 are xylem sap-specific proteins that are required for full *V. dahliae* pathogenicity on tomato. These factors play a role during the colonization phase and represent potential targets for new control strategies for Verticillium wilt.

## Introduction

Vascular wilts caused by *Verticillium* spp. are widespread and destructive plant diseases, resulting in enormous economic losses. Haploid *Verticillium dahliae*, the economically most important representative of the genus, infects over 200 plant species worldwide [1, 2]. In contrast, the allodiploid *Verticillium longisporum* has a narrow host range that comprises primarily *Brassicaceae*. During the last several decades, increasing cultivation of the oilseed rape *Brassica napus* revealed *V. longisporum* as one of the most devastating pathogens of oilseed rape [3]. *Verticillium* spp. enter the plant through the root, where the fungus then grows both inter- and intracellular in the root cortex towards the central cylinder and finally colonizes the xylem vessels [4, 5]. The transpiration stream plays an essential role in supplying water and mineral salts to the aerial tissue of plants [6]. The xylem sap is a nutrient-poor environment with plant defense proteins, hormones and low concentrations of amino acids and sugars [7, 8]. This makes it a very unique environment, which *Verticillium* spp. exploit for growth and systematic distribution in the host plant [7, 9]. Factors that contribute to adaptation to the unbalanced amino acid supply include the chorismate synthase encoding gene *VlARO2* and the cross-pathway transcription factor *CPC1* [7, 10]. The pathogen requires distinct sets of enzymes during different steps in plant colonization including carbohydrate active enzymes (CAZys) and peptidases, as well as small secreted proteins, to establish an infection and overcome the immune response of the plant. Several extracellular proteins including polygalacturonases, pectate lyases, xylanases or lipases presumably contribute to virulence during pathogen-host interactions [11–15].

The plant immune response depends in part on transmembrane receptor proteins termed pattern recognition receptors (PRRs). Cell surface localized PRRs recognize conserved microbial molecules and structural motifs designated as pathogen-associated molecular patterns (PAMPs); examples include the fungal cell-wall polymer chitin [16]. PAMP perception elicits a basal defense response which halts colonization by non-adapted pathogens and results in PAMP-triggered immunity (PTI). Host-adapted pathogens circumvent PTI by secretion of specific effector proteins as virulence factors for different phases of the infection cycle [17]. These secreted effectors may act passively or actively to combat plant defense responses [18].

Well known examples of fungal effectors include the Avr4 and Ecp6 effectors from the leaf mold fungus *Cladosporum fulvum* that bind to chitin oligosaccharides via a carbohydrate-binding module (CBM) or LysM domain, respectively [18–21]. Similarly, a chitin scavenging function has also been assigned to Cp1 in *V. dahliae* strain XH-8. *CP1* knockout mutants were affected in cotton virulence [22]. This chitin protection leads to the suppression of the PTI of the plant and shields the fungal cell wall from plant chitinases that hydrolyze chitin [18–21]. Other fungal effectors such as metalloproteases possess enzymatic activity and are able to truncate plant chitinases that attack the fungal cell wall [23, 24]. Toxins provide another means for pathogens to attack plant hosts. For example, necrosis and ethylene inducing-like proteins (NLP) induce immune responses and cell death in host tissues and are conserved among fungi including *Verticillium* spp. [25, 26]. *V. dahliae* isolates encode up to eight NLP homologs [26, 27] whereas most other fungi generally only possess up to three NLPs [25]. Only Nlp1 and Nlp2 show cytotoxic activity in *Nicotiana benthamiana* leaves and play strain- and host-specific roles in *V. dahliae* virulence [26, 27]. Nlp1 and Nlp2 are required for *V. dahliae* JR2 pathogenicity on tomato and *A. thaliana* [26] whereas the corresponding proteins in *V. dahliae* V592 did not alter virulence on cotton [27]. Plant pathogens additionally require a set of carbohydrate-active enzymes (CAZys) that facilitate the breakdown of the plant cell wall [28]. The genomes of *Verticillium* species encode a greater number of cell wall-degrading enzymes with a strikingly high repertoire of pectin-degrading enzymes compared to the secreted proteins of other plant pathogens [18, 29].

As the fungus lives in the vascular tissues during most of its life cycle in the plant, further knowledge about specific secretion responses would enable a better understanding of fungal-plant interactions during the infection process. Once the fungus resides inside the plant, it is inaccessible for pesticides and therefore the management of Verticillium wilt is very challenging. The most effective and widely used soil fumigants, methyl bromide or metam sodium, are used for high valuable crops, but are not profitable for all crops. Furthermore, these and other banned fungicides, are associated with environmental issues [1, 30]. Therefore, an indispensable approach for protection is to use resistant plant varieties, but these are not available in most crops. The selection pressure on fungal strains to quickly overcome genetic resistances of the plant makes it even more difficult to develop new resistant varieties [1, 18]. Consequently, an increased understanding of the infection process for *Verticillium* spp. is necessary to identify new approaches for disease control.

Until now, it is not known how the effector repertoire of *Verticillium* spp. is expressed once the pathogen enters the plant. Analysis of *Verticillium* strains in their vascular environment is technically demanding, and a large number of plants are required to harvest sufficient amounts of xylem sap. Proteomic approaches can be fruitful because comparison of the intracellular fungal proteome in diluted xylem sap and pectin-rich medium resulted in the identification of a disease-related catalase peroxidase, which was only up-regulated in the presence of xylem sap and not in the presence of pectin [31].

In this study, we extended the comparative analysis with rapeseed xylem sap and focused on the fungal secretome. *V. longisporum* secreted proteins that were derived from cultivation in different growth media were identified by a proteomic approach and the protein patterns induced by different environments were compared. Our goal was to obtain a more comprehensive overview of the secreted factors of *V. longisporum* in response to different substances in its environment that putatively reflect different stages of the infection. We analyzed the exoproteomes of *V. longisporum* on a broad range of media from water to minimal and complete media. As an additional condition, we applied simulated xylem medium (SXM), which is rich in pectin and which was originally developed to mimic the natural plant growth environment, the xylem sap [32]. All exoproteomes were compared to fungal cultures grown in extracted xylem sap of the *V. longisporum* host oilseed rape *B. napus*.

Our results demonstrate that *V*. *longisporum* is able to distinguish between the different environments to express different secretome patterns. The pectin-rich medium and xylem sap each triggered distinct protein patterns in comparison to all other tested media. The fungal response to growth in the pectin-rich medium and xylem sap consists of a shared core exoproteome and an additional group of uniquely secreted proteins. A small number of proteins are specifically expressed in xylem sap including CAZys and other potential virulence factors. Of the factors that were specifically enriched in xylem sap, the NLPs, Nlp2 and Nlp3 proteins, we demonstrate contributions to plant pathogenicity as virulence factors.

## Results

### Xylem sap and pectin-rich SXM trigger specific exoproteomic patterns compared to other growth media

*V. longisporum* is a rapeseed pathogenic fungus that is able to grow on a variety of different substrates and colonizes the xylem vessels of plants. This requires the adaptation of the fungus to changing nutrient conditions and other biotic and abiotic factors. We examined how different growth media affected the exoproteome of *V*. *longisporum* with a specific focus on identification of distinct patterns triggered by different plant-related contents. For all experiments, *V. longisporum* was precultured in liquid potato dextrose medium (PDM) to ensure an equal initial growth state prior to the media-specific induction of secretion. The proteins of the *V. longisporum* culture supernatants from different media were precipitated and separated by one-dimensional SDS-PAGE thereby resulting in several different patterns in colloidal Coomassie stained gels (Fig 1A). The defined media conditions corresponded to different levels of complexity including nutrient-free water, water with glucose as carbon source, and a more complex nitrogen-rich medium (YNB: Yeast Nitrogen Base). Several plant-related media were included because the natural habitat of the pathogen *V. longisporum* is inside host plants. These media included the nutrient-limited sucrose medium CDM (Czapek-Dox medium), which was either supplemented with 7% of *B. napus* xylem sap or plant proteins, vegetable juice (V8 juice) and the pectin-rich SXM, which was developed to mimic fungal growth conditions in plants *in vitro* [32, 33]. Finally, we also used extracted xylem sap from the rapeseed plant *B. napus*, corresponding to the natural habitat of the fungus.

**Fig 1.**
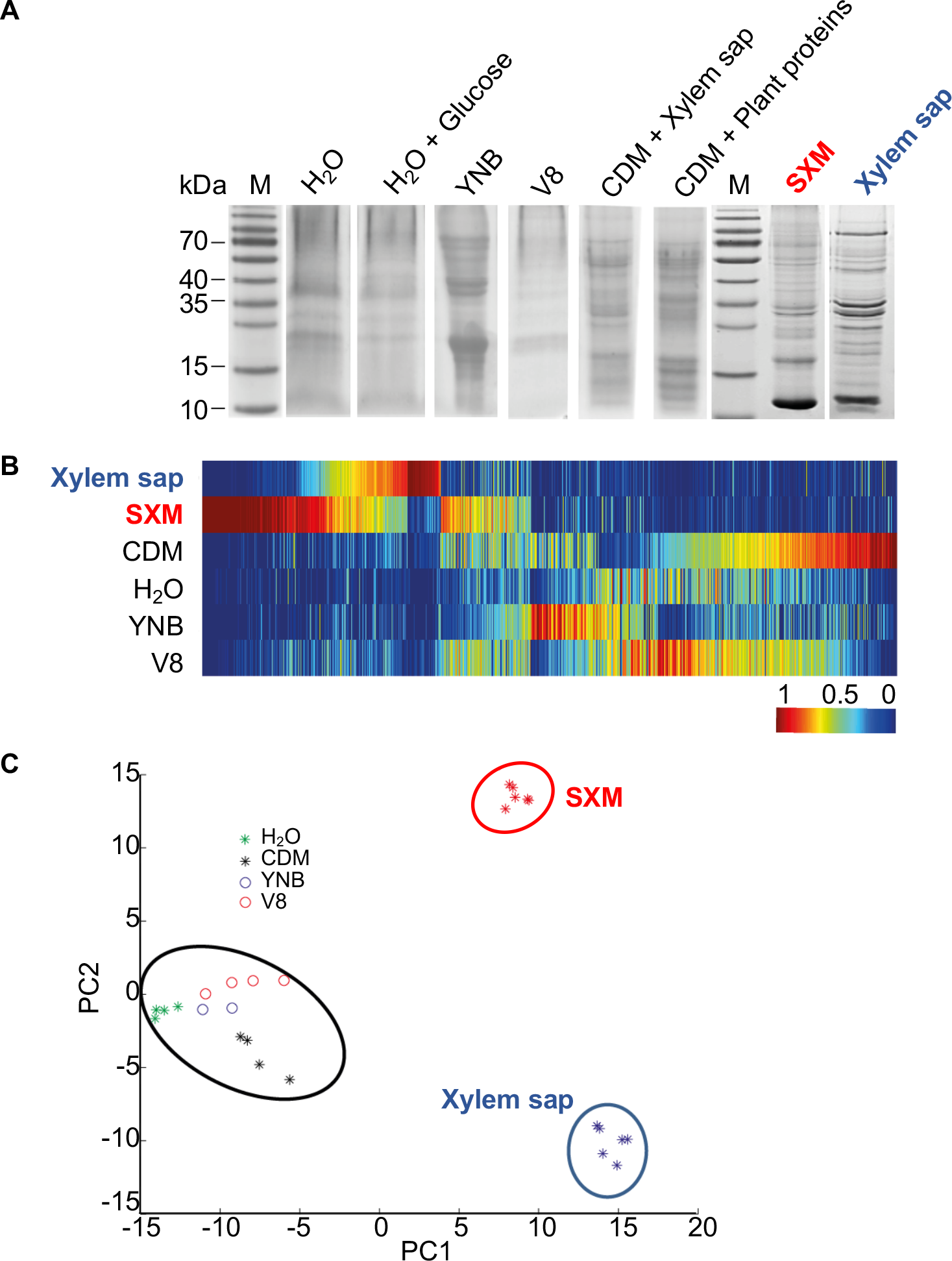
Exoproteome signatures of *V. longisporum* in different growth media. *V. longisporum* 43 was cultivated in complete medium (PDM) for four days before the sedimented mycelia and spores were dissolved in different media (dH_2_O, dH_2_O + 0.1% glucose, yeast nitrogen base (YNB), vegetable juice (V8), minimal medium (CDM) supplemented with 7% xylem sap or plant proteins, simulated xylem medium (SXM) and extracted xylem sap from *B. napus*) and cultivated for four more days. **(A)** Precipitated proteins from the supernatants were separated by one-dimensional SDS-PAGE. The colloidal Coomassie stained exoproteome samples on gel displayed a strong variation between the protein band distributions of all different culture conditions. Lanes M represent the molecular weight marker. Complete lanes of the *V. longisporum* exoproteomes were subjected to tryptical protein digestion and the resulting peptides were analyzed by LC MS/MS. **(B)** Clustering analysis of protein abundances (spectral counts) was facilitated by the software MarVis and is visualized as one-dimensional self-organizing maps. Rows represent the compared growth conditions. The spectral counts were normalized and color-coded according to the indicated scale. Red indicates increased, dark blue no spectral counts. **(C)** The principle component analysis plot of the exoproteomes based on the spectral counts was performed with MarVis-software [67]. Each dot represents one biological replicate (one independent culture). The compared exoproteome signatures cluster in three groups: a first cluster is formed by all xylem sap culture samples (blue circle); the second cluster contains all SXM culture samples (red circle) and the third cluster consists of all other samples (black circle).

To obtain a more comprehensive analysis, complete lanes of the gels with the respective exoproteomes were fractionally subjected to tryptic protein digestion and the resulting peptides were analyzed by LC MS/MS. The obtained raw data were channeled through a bioinformatics pipeline based on Proteome Discoverer Software 1.3™ (Thermo Scientific) and an in-house genome-wide protein sequence database of *V. longisporum*. The received spectral counts were compared on single secreted protein level by color-coded one-dimensional self-organizing maps (Fig 1B). These revealed that proteins that were strongly enriched in xylem sap or SXM were not enriched in any other condition. Differences in the exoproteome signatures are also illustrated by sample clustering in a principle component analysis plot (Fig 1C). Exoproteomes of *V. longisporum* derived from very diverse media including nutrient-free water, V8 juice, CDM or YNB medium show a similar pattern. Supplementation of CDM with *B. napus* plant proteins or xylem sap with a final concentration of 7% did not result in a different exoproteome pattern, neither did glucose supplementation to water. Therefore, the respective results for these conditions were combined together. In contrast, proteins secreted in pectin-rich SXM or xylem sap each showed a distinct pattern in comparison to the other media conditions. These patterns representing the latter exoproteomes are similar in some features as the clusters lie close to each other on the x-axis, although some differences are present as analyzed further below.

Overall, our analysis illustrates that the fungus has the potential to form a general secretome response to non-plant related environments and, in addition, a similar, but more specialized response to plant-related substances (Fig 1C).

### Pectin-rich medium and xylem sap elicit distinct *V. longisporum* exoproteome responses

Similarities and differences of the specific *V*. *longisporum* exoproteome responses in the pectin-rich SXM compared to xylem sap were analyzed in more detail in a large-scale experimental set up. The protein patterns of six biological replicates of each cultivation condition were compared. The proteins were precipitated from the culture supernatants, subjected to LC MS/MS and analyzed. The data set was filtered with a statistical workflow using MarVis-Suite [34]. S1 Table shows the identities of the 445 proteins from an in-house database with their protein sequences and details on their abundance as measured by identified peptides. The list is sorted according to the most specifically enriched proteins in xylem sap (green) and SXM (red), respectively. Proteins that are not considered as specifically enriched are listed at the bottom of the table and belong to the core exoproteome.

Clustering analysis of spectral count data of the 445 proteins with MarVis-Suite was visualized as a one-dimensional self-organizing map (Fig 2A). Upper and lower rows represent the two growth conditions, SXM and xylem sap, and each column corresponds to one identified protein. The spectral counts were normalized and color-coded according to the indicated scale where red columns indicate increased and dark blue columns no spectral counts. A set of proteins, which have a stable abundance in both conditions is considered as the shared core exoproteome. Proteins that showed different peptide counts in the two growth conditions were considered as differentially enriched (Fig 2A, ‘enriched in Xylem sap’ and ‘enriched in SXM’, respectively).

**Fig 2.**
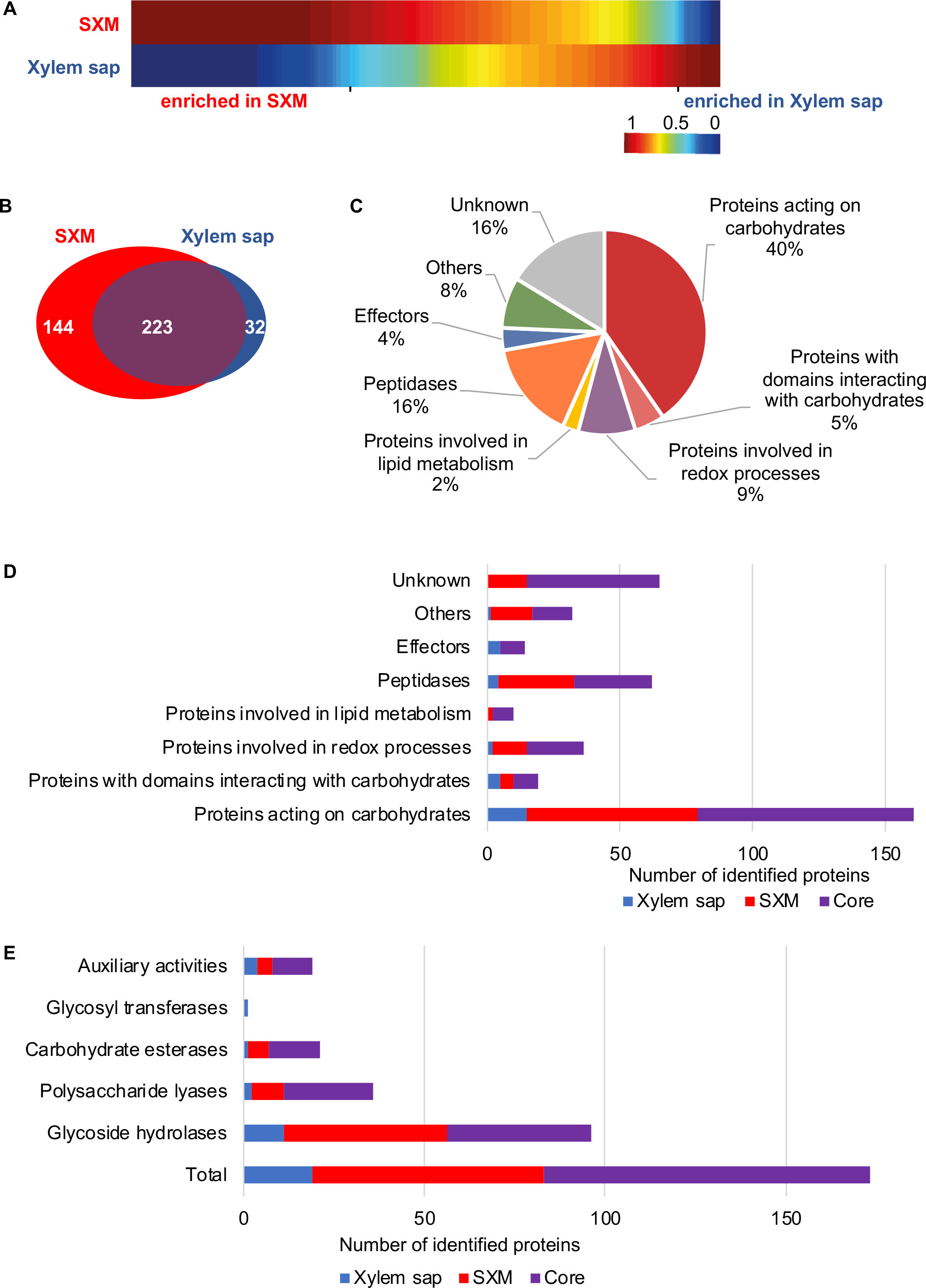
Comparison of exoproteome signatures for *V. longisporum* growth in xylem sap and pectin-rich simulated xylem medium. Complete lanes of the SDS-gel with samples of *V. longisporum* simulated xylem medium (SXM) and xylem sap derived exoproteomes were subjected to tryptical protein digestion and the resulting peptides were analyzed by LC MS/MS. The obtained raw data were searched with Proteome Discoverer Software 1.3™ against a draft genome-wide protein sequence database of *V. longisporum*. Lists of identified proteins were semi-quantitatively processed using MarVis-Suite [34]. Single proteins with identifiers are found in S1 Table. **(A)** Clustering analysis of protein abundances (spectral counts) is visualized as one-dimensional self-organizing maps, which was facilitated by the software MarVis. Upper and lower rows represent the two compared growth conditions xylem sap and pectin-rich SXM, respectively. Each column corresponds to spectral counts of one identified protein. The spectral counts were normalized and color-coded according to the indicated scale. Red indicates increased, dark blue no spectral counts. **(B-E)** BLAST searches of identified *V. longisporum* proteins in **A** against the *V. dahliae* JR2 and the *V. alfalfae* VaMs.102 proteomes from Ensembl Fungi [35] were conducted and functional analysis is based on the *V. dahliae* JR2 or *V. alfalfae* VaMs.102 protein sequences. **(B)** The Venn diagram displays the number of proteins specifically enriched in xylem sap (blue) and proteins enriched in SXM (red). All proteins below the statistical threshold form the core exoproteome that is similarly enriched in both media (violet). **(C)** The cake diagram shows the functional classification of the 399 identified secreted proteins into main protein groups according to their predicted domains. **(D)** The functional groups are presented with the number of identified proteins in the different cultivation environments. **(E)** Classification by CAZy modules is shown for each cultivation type.

*A. V. longisporum* is an allodiploid organism derived from two parental strains, and *V. longisporum* 43 used in this study is a result of an A1xD1 hybridization event. A1 and D1 are described as so far unknown haploid *Verticillium* species, of which D1 is closely related to *V. dahliae* and A1 is distantly related to *V. alfalfae* [2]. Most genes are encoded in two copies, reflecting the two isogenes of both parental lineages. BLAST searches against the *V. dahliae* JR2 and the *V. alfalfae* VaMs.102 proteomes from Ensembl Fungi [35] were conducted. As a consequence, two isogene products were detected for most identified proteins.

The list of proteins was further and thoroughly analyzed manually to functionally classify the candidates. As the Ensembl Fungi annotations are more robust, further analyses are based on the *V. dahliae* JR2 protein sequences except for candidates with no corresponding hit in *V. dahliae*. Here, the protein sequences for further analyses were retrieved from the *V. alfalfae* VaMs.102 proteome [35]. Putative functions of robust annotated proteins were addressed with InterProScan [36], the CAZy database (http://www.cazy.org) and dbCAN2 [37]. All details are given in S2 Table. The Venn diagram in Fig 2B displays the 399 candidates with robust annotations in different groups. Protein extracts from pectin-rich SXM and xylem sap share a core exoproteome of 223 proteins with a similar abundance in both media, but each also induced the secretion of distinct exoprotein patterns. SXM cultivation resulted with 144 secreted proteins in a four-fold higher number of secreted proteins specifically enriched in comparison to xylem sap, where the peptide count of 32 proteins was specifically increased (Fig 2B).

These results show that SXM, which is used to simulate xylem sap *in vitro,* and xylem sap of *B. napus* induce distinct secretion responses with different facets. This indicates that the fungus is able to fine-tune its secretion responses.

### *V. longisporum* secretes a broad arsenal of substrate-degrading enzymes in pectin-rich medium and in xylem sap

All 399 identified secreted proteins were classified into functional groups according to the predicted domains (S2 Table, Fig 2C). For 65 identified proteins, classified as hypothetical gene products, no information about structural domains or putative functions could be found (‘Unknown’). ‘Proteins involved in lipid metabolism’, ‘Effectors’ and smaller groups combined as ‘Others’ represent minor groups. More proteins were sorted to ‘Proteins involved in redox processes’ (9%) or ‘Peptidases’ (16%) whereas the functional classification revealed an overrepresentation of proteins involved in carbohydrate metabolism or catabolism with around 40% of proteins acting on carbohydrates and another 5% of proteins with domains interacting with carbohydrates (Fig 2C, S3 Table).

Further analysis of proteins from functional groups regarding the induction by different media showed that pectin-rich medium predominantly triggered the secretion of carbohydrate-degrading enzymes, but also peptidases and redox enzymes (Fig 2D, S3 Table). No effectors were found in the SXM-specific exoproteome. Cultivation in xylem sap triggered the unique secretion of carbohydrate-degrading enzymes, effectors, peptidases and redox enzymes though the total number of enriched proteins is significantly smaller compared to the SXM-specific secreted proteins.

The majority of identified secreted proteins comprise the carbohydrate-active enzymes, which contain protein motifs that have been classified into sequence-related families of CAZy modules [38]. Within the group of all secreted proteins we identified 96 glycoside hydrolases (GHs), 36 polysaccharide lyases (PLs), 21 carbohydrate esterases (CEs), one glycosyltransferase (GT), 19 auxiliary activities (AAs). Of these CAZys, 21 proteins additionally possess non-catalytic, carbohydrate-binding modules (CBMs) (Fig 2E, S4 Table). In the proteins from the pectin-rich medium condition, the CAZys are highly represented with 64 proteins whereas in xylem sap only 19 CAZys are specifically enriched and 90 proteins belong to the core exoproteome (S4 Table). The core exoproteome exhibits an overrepresentation of CAZy families with 32 proteins acting on pectin, including members of family GH28 (five proteins), PL1 (16 proteins), PL3 (seven proteins) and CE8 (four proteins). Additionally, the SXM-specific and most enriched CAZy families comprise 20 pectin-degrading enzymes (families GH28, PL1 and CE12 with ten, six and four proteins, respectively, S4 Table). Only a few CAZys were specifically enriched in the xylem sap condition, and these were distributed in different families.

Overall, we found that *B. napus* xylem sap and pectin-rich SXM, employed as plant-related culture environments, predominantly induced the secretion of carbohydrate-degrading enzymes. Compared to the rapeseed xylem sap condition, SXM triggered an additional set of CAZys that were specifically enriched after cultivation in this medium.

### Xylem sap triggers the secretion of potential and known *Verticillium* effectors

Compared to the pectin-rich SXM, *V. longisporum* formed a more specific secretion response in xylem sap with only 32 proteins that are uniquely enriched. This indicates that the fungus can distinguish between xylem sap and the presence of other plant material and accordingly fine-tunes its protein secretion. Furthermore, the proteins secreted in the host xylem sap might be specifically important during plant colonization. Table 1 displays the xylem sap-specific proteins. The corresponding isogenes are paired up and the best hit in *V. dahliae* JR2 is given, except for the *V. alfalfae* specific proteins that were searched against the *V. alfalfae* VaMs.102 proteome (Ensembl Fungi). Within the identified groups, proteins were ranked according to the quotient of peptide counts identified in xylem sap (XyS) by the number detected in the pectin-rich SXM. Displayed peptide counts were averaged from 6 biological replicates. If the number was below 1, it was calculated as 0 and the quotient was given as the average XyS peptide counts (>). The 32 xylem sap-specific proteins comprise 15 ‘Proteins acting on carbohydrates’, five ‘Proteins with domains interacting with carbohydrates’, five ‘Effectors’, four ‘Peptidases’, two ‘Proteins involved in redox processes’ and one with a ubiquitin binding domain that was grouped as ‘Other’ (Table 1).

**Table 1.** Xylem sap-specific exoproteins of V. longisporum.

|  | Name | VI43 identifier | Best Hit *V. dahliae* JR2 | Protein domain | Peptide counts |  | XyS/<br>SXM | s/l<br>ratio |
| --- | --- | --- | --- | --- | --- | --- | --- | --- |
|  |  |  |  |  | SXM | XyS |  |  |
| Proteins acting on carbohydrates | **Gla1** | vl43-au16.g17458.t1 | VDAG_JR2_Ch8g11020a | Glycoside hydrolase 15, Carbohydrate binding module family 20 | 0 | 8.2 | **>8.2** | 0.76 |
|  |  | vl43-au16.g13024.t1 |  |  | 0 | 3.8 | **>3.8** | 0.62 |
|  | **Amy1** | vl43-au16.g6892.t1 | VDAG_JR2_Ch7g03330a | Glycoside hydrolase 13 | 1.5 | 12 | **8** | 0.68 |
|  |  | vl43-au16.g9360.t1 | *V. alfalfae*: VDBG_05827 | Glycosyl hydrolases 134 | 0 | 7.5 | **>7.5** | 0.77 |
|  |  | vl43-au16.g6746.t1 | VDAG_JR2_Ch3g11080a | Glycoside hydrolase 43 | 1.2 | 7.8 | **6.5** | 0.68 |
|  |  | vl43-au16.g12522.t1 |  |  | 1 | 5.8 | **5.8** | 0.61 |
|  |  | vl43-au16.g12596.t1 | VDAG_JR2_Ch6g04040a | Glycosyl hydrolases 11 | 0.8 | 5.2 | **>5.2** | 0.55 |
|  |  | vl43-au16.g15027.t1 | VDAG_JR2_Ch6g09340a | Polysaccharide lyase 6 | 2.2 | 11.2 | **5.1** | 0.57 |
|  |  | vl43-au16.g5309.t1 |  |  | 2.3 | 10.8 | **4.7** | 0.59 |
|  |  | vl43-au16.g207.t1 | VDAG_JR2_Ch1g06940a | Glycoside hydrolase 16 | 2.2 | 9 | **4.1** | 0.56 |
|  |  | vl43-au16.g14986.t1 |  |  | 2.3 | 9.5 | **4.1** | 0.55 |
|  |  | vl43-au16.g15947.t1 | VDAG_JR2_Ch3g00250a | Glycoside hydrolase 131 | 1 | 4 | **4** | 0.43 |
|  |  | vl43-au16.g9097.t1 | *V. alfalfae*: VDBG_03110 | Glycoside hydrolase 12 | 1 | 2.8 | **2.8** | 0.38 |
|  |  | vl43-au16.g16765.t1 | VDAG_JR2_Ch1g04240a | Glycosyl transferase 1 | 0.3 | 1.7 | **>1.7** | 0.49 |
|  |  | vl43-au16.g9025.t1 | VDAG_JR2_Ch7g01210a | Carbohydrate esterase 1, Carbohydrate binding module family 1 | 0.2 | 1.3 | **>1.3** | 0.49 |
| Proteins with domains interacting with carbohydrates | **Cbd1** | vl43-au16.g11636.t1 | VDAG_JR2_Ch4g04440a | Auxiliary activity 13, Carbohydrate binding module family 20 | 1.7 | 13.8 | **8.1** | 0.71 |
|  |  | vl43-au16.g3945.t1 |  |  | 1.3 | 6.8 | **5.2** | 0.55 |
|  |  | vl43-au16.g5519.t1 | VDAG_JR2_Ch6g09790a | Auxiliary activity 9 | 2 | 8.2 | **4.1** | 0.51 |
|  |  | vl43-au16.g20506.t1 |  |  | 2.3 | 8.8 | **3.8** | 0.46 |
|  |  | vl43-au16.g11273.t1 | VDAG_JR2_Ch6g05220a | WSC Carbohydrate binding domain | 3.2 | 11.7 | **3.7** | 0.46 |
| Effectors | **Nlp3** | vl43-au16.g9727.t1 | VDAG_JR2_Ch4g05950a | Necrosis inducing protein | 0.3 | 8.3 | **>8.3** | 0.84 |
|  |  | vl43-au16.g96.t1 |  |  | 0.2 | 6.3 | **>6.3** | 0.81 |
|  | **Nlp2** | vl43-au16.g3884.t1 | VDAG_JR2_Ch2g05460a | Necrosis inducing protein | 1 | 4.8 | **4.8** | 0.61 |
|  |  | vl43-au16.g12566.t1 |  |  | 1.8 | 6.5 | **3.6** | 0.47 |
|  | **Cp2** | vl43-au16.g16459.t1 | VDAG_JR2_Ch2g07000a | Cerato-platanin | 0.3 | 1.8 | **>1.8** | 0.41 |
| Peptidases | **Mep2** | vl43-au16.g19470.t1 | VDAG_JR2_Ch1g21900a | Peptidase M43 | 1 | 8.8 | **8.8** | 0.57 |
|  |  | vl43-au16.g11262.t1 |  |  | 1.3 | 8.7 | **6.7** | 0.55 |
|  | **Mep1** | vl43-au16.g19320.t1 | VDAG_JR2_Ch8g09760a | FTP domain, Peptidase M36, fungalsin | 1.7 | 14 | **8.2** | 0.69 |
|  |  | vl43-au16.g14388.t1 |  |  | 1.8 | 13.5 | **7.5** | 0.65 |
| Proteins involved in redox processes |  | vl43-au16.g2684.t1 | VDAG_JR2_Ch8g10630a | FAD-binding domain | 6 | 13.2 | **2.2** | 0.33 |
|  |  | vl43-au16.g13340.t1 |  |  | 8 | 15.3 | **1.9** | 0.34 |
| Other |  | vl43-au16.g18734.t1 | VDAG_JR2_Ch1g04640a | Ubiquitin 3 binding protein But2, C-terminal | 9.7 | 18.8 | **1.9** | 0.37 |

VI43 = *V. longisporum* 43; SXM = simulated xylem medium; XyS = xylem sap; s/l = signal-to-level; FTP = fungalsin/thermolysin propeptide; WSC = putative carbohydrate-binding domain

Several potential virulence factors were identified in the xylem sap-specific response set. Proteins involved in the degradation of carbohydrates are known to contribute to *V. dahliae* pathogenicity [39, 40]. The five proteins comprising domains of already characterized *V. dahliae* effectors incorporate either necrosis-inducing protein (NPP1, also necrosis-inducing *Phytophthora* protein) or cerato-platanin (CP) domains. NPP1 domains are characteristic for necrosis and ethylene inducing-like proteins (NLP) of which *Verticillium* spp. contain up to eight members [26]. Nlp1 and Nlp2 were previously shown to differentially contribute to *V. dahliae* pathogenicity on different hosts [26, 27, 41]. Our secretome approach identified four isogene products corresponding to two NLPs, Nlp2 and Nlp3. Of the CP domain-containing proteins, one isogene assigned to Cp2 was identified as specifically enriched in xylem sap.

*V. dahliae* possesses two CP proteins of which Cp1 affects virulence on cotton [22]. Additionally, four isogenes of two metallopeptidases were found in the xylem sap-specific secretome. Metalloproteases are able to truncate host defense proteins such as chitinases and therefore have the potential to act as virulence factors [23]. These findings show that *V. longisporum* finetunes its protein secretion response in the host xylem sap, which include known and potential effectors important for plant colonization or infection.

### Xylem sap-specific secreted proteins are dispensable for *V. dahliae ex planta* development

*V. longisporum* formed a specific secretion response in xylem sap compared to other media showing that the fungus can distinguish between xylem sap and the presence or absence of other plant material. To investigate whether the xylem sap-specific proteins play a major role in fungal colonization of the plant, the top candidates of the protein groups that have been shown to play critical roles in plant colonization [5, 22, 26] were analyzed in this study.

Chosen proteins are highlighted in Table 1 with a bold given name. The follow up genetic analyses of these proteins were conducted with *V. dahliae* JR2 because it can be more easily manipulated genetically compared with the allodiploid *V. longisporum* strain. All candidates from the group “Effectors” were included in the follow up experiments. These comprise two NLPs, Nlp2 and Nlp3, and the Cp2 protein. The *V. dahliae* JR2 homolog of *CP1* was tested as well. Cp1 is a virulence factor of *V. dahliae* strain XH-8 in infections on cotton [22]. The peptidases that were identified as specifically enriched in xylem sap were named Mep1 and Mep2 and were both included in our genetic analyses. Of the largest group, proteins involved in carbohydrate degradation, the three most highly abundant candidates, the glucoamylase Gla1, the carbohydrate-binding module containing protein Cbd1, and the *α*-amylase Amy1, were further investigated.

For the construction of the corresponding deletion mutants, the open reading frame (ORF) was replaced with either the nourseothricin or hygromycin resistance cassette under control of the constitutively active *gpdA* promoter and *trpC* terminator. Correct integration of the deletion cassette was verified by Southern hybridization (S1-S4 Figs). To investigate a putative combined effect of proteins in similar groups, the following double deletion strains were constructed as well: *NLP2*/*NLP3*, *MEP1/MEP2* and *CP1/CP2*. Ectopic complementation strains were also constructed for the *CP1* and *CP2* deletion mutants. Phenotypical analysis of all strains revealed no alteration compared to *V. dahliae* JR2 wildtype (WT) growth and development on solid agar plates such as minimal and complete medium, and simulated xylem medium. Additionally, the strains were tested for the involvement in stress responses with at least one stressor tested for each strain. The stress inducing agent was added to minimal medium. The cell wall perturbing agents SDS and ethanol or the oxidative stressor hydrogen peroxide were used. All single deletion, double deletion and complementation strains exhibited a similar morphological development to *V. dahliae* WT, which is exemplified by growth on SXM (Fig 3).

**Fig 3.**
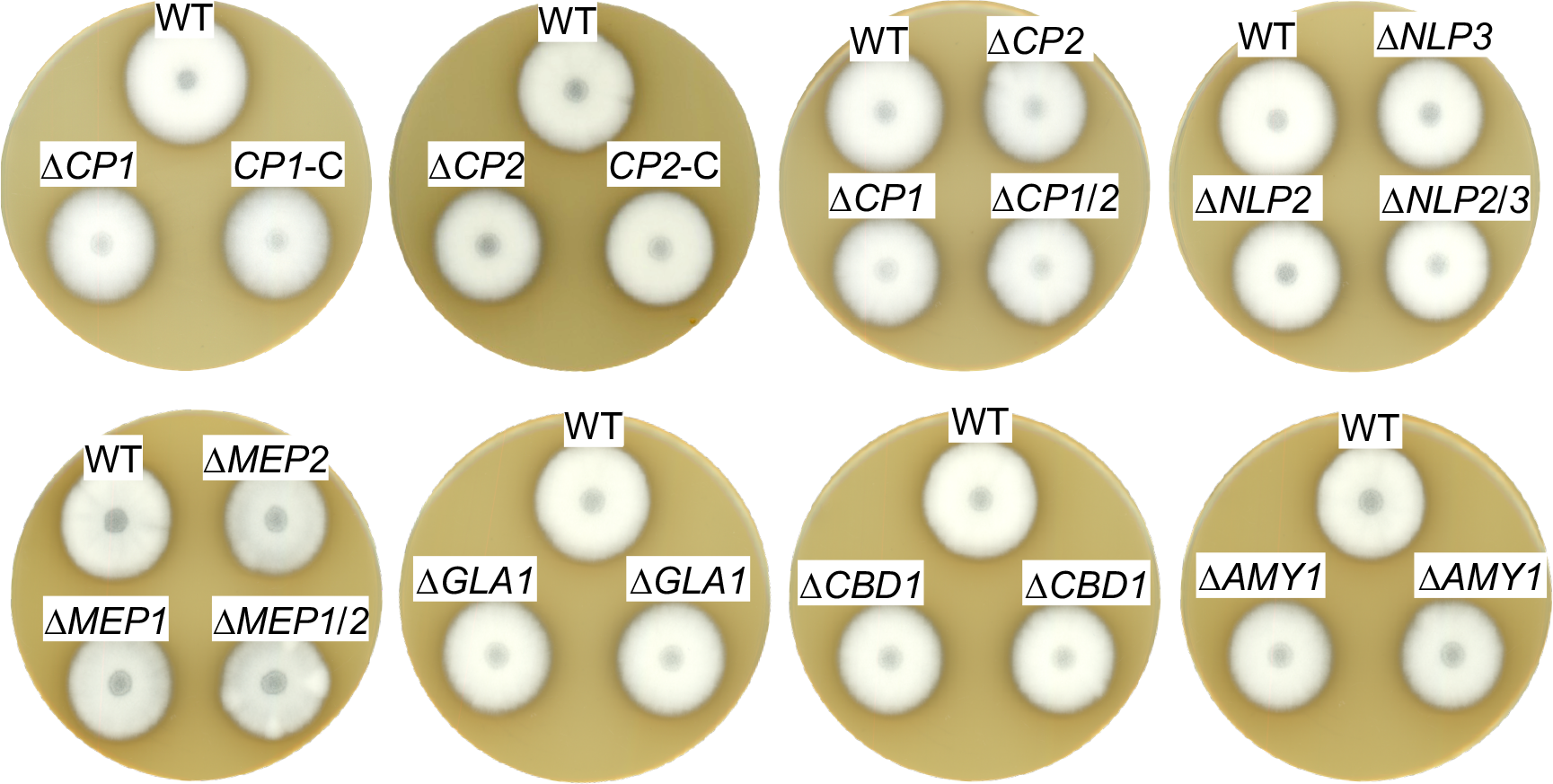
Exoproteins specifically secreted in xylem sap are dispensable for *V. dahliae ex planta* phenotype. The same number of spores of *V. dahliae* JR2 wildtype (WT) and indicated deletion mutant (Δ*CP1*, ΔCP2, Δ*CP1/2*, Δ*NLP2*, Δ*NLP3*, Δ*NLP2/3*, Δ*MEP1*, Δ*MEP2*, Δ*MEP1/2*, Δ*GLA1*, Δ*CBD1*, Δ*AMY1*) and complementation (*CP1*-C, *CP2*-C) strains were point inoculated on simulated xylem medium (SXM) plates and incubated at 25°C for 10 days. For Δ*AMY1*, Δ*GLA1* and Δ*CBD1* mutants two transformants were spotted. Top-view scans of the colonies show a similar phenotype of all strains.

Overall, these results suggest that Cp1, Cp2, Nlp2, Nlp3, Mep1, Mep2, Gla1, Cbd1 and Amy1, that were found to be enriched specifically in xylem sap cultures, are dispensable for vegetative growth, development and stress response of *V. dahliae*.

### Xylem sap-specific CAZys, metalloproteases and cerato-platanin proteins are dispensable for *V. dahliae* JR2 pathogenicity in tomato infections

The functions of the proteins that were specifically enriched after cultivation in xylem sap would be predicted to be important in the interaction with plant substrates in the host xylem sap. Therefore, all *V. dahliae* single and double deletion strains were tested for their virulence on tomato. Ten-day-old tomato seedlings were root-inoculated with the indicated mutant strains, and plants treated with demineralized water were used as mock controls. Disease symptoms were measured three weeks after inoculation and included the height of the plant, the longest leaf length and the fresh weight of the aerial part of the plant. The stack diagrams display the percentage of plants exhibiting the respective symptoms (Figs 4, 5). We found that plants infected with *GLA1, CBD1, AMY1, MEP1, MEP2, MEP1/MEP2, CP1, CP2, CP1/CP2* deletion strains or *CP1* or *CP2* complementation strains showed similar disease symptoms as the WT-infected plants. That is, all fungal infections resulted in a similar stunting phenotype as WT colonization, and plant defense reactions were observed by the discoloration of the hypocotyls in all infected plants.

**Fig 4.**
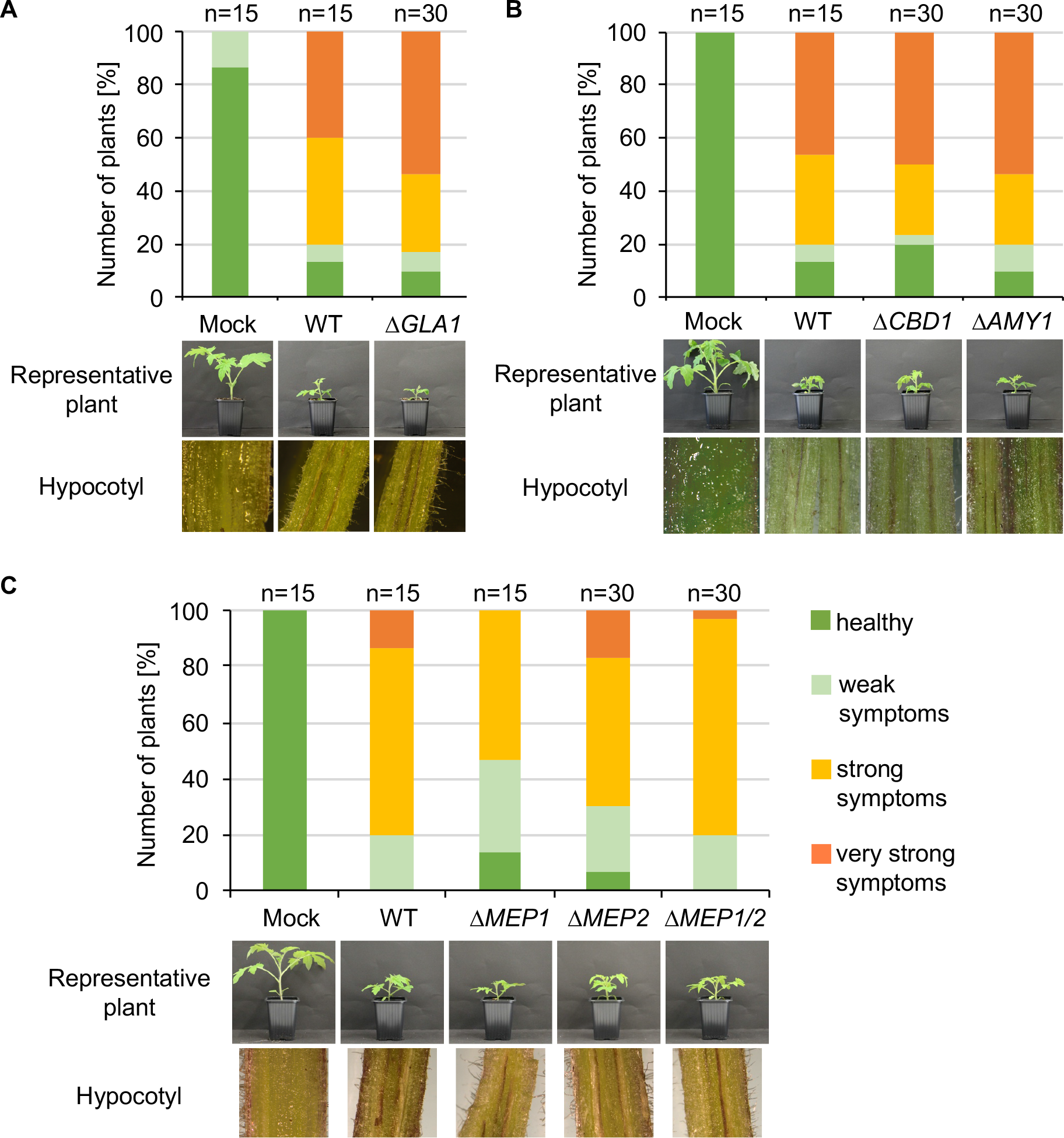
CAZys and metalloproteases specifically secreted in xylem sap are dispensable for *V. dahliae* pathogenicity on tomato. Ten-day-old tomato seedlings were root-infected with spores of *V. dahliae* JR2 (WT) and the indicated single and double deletion strains (Δ*GLA1*, Δ*CBD1*, Δ*AMY1, ΔMEP1, ΔMEP2, ΔMEP1/2*). Uninfected plants (mock) served as control. The disease index was assessed after 21 days of growth in the climate chamber under 16h:8h light:dark at 22-25°C and includes the height of the plant, length of the 2^nd^ true leaf and weight of the plant. Representative plants and discolorations of the hypocotyls are shown for each infection. The number (n) of plants is shown for each fungal strain or mock treatment. **(A)** Infections with *GLA1* (glycoamylase) deletion strains resulted in the same stunting phenotype as WT infections. **(B)** Strains with deletion of *CBD1* (carbohydrate-binding domain) or *AMY1* (amylase) resulted in a WT-like induction of disease symptoms. **(C)** Absence of metalloproteases Mep1 and Mep2 resulted in WT-like *V. dahliae* pathogenicity on tomato.

**Fig 5.**
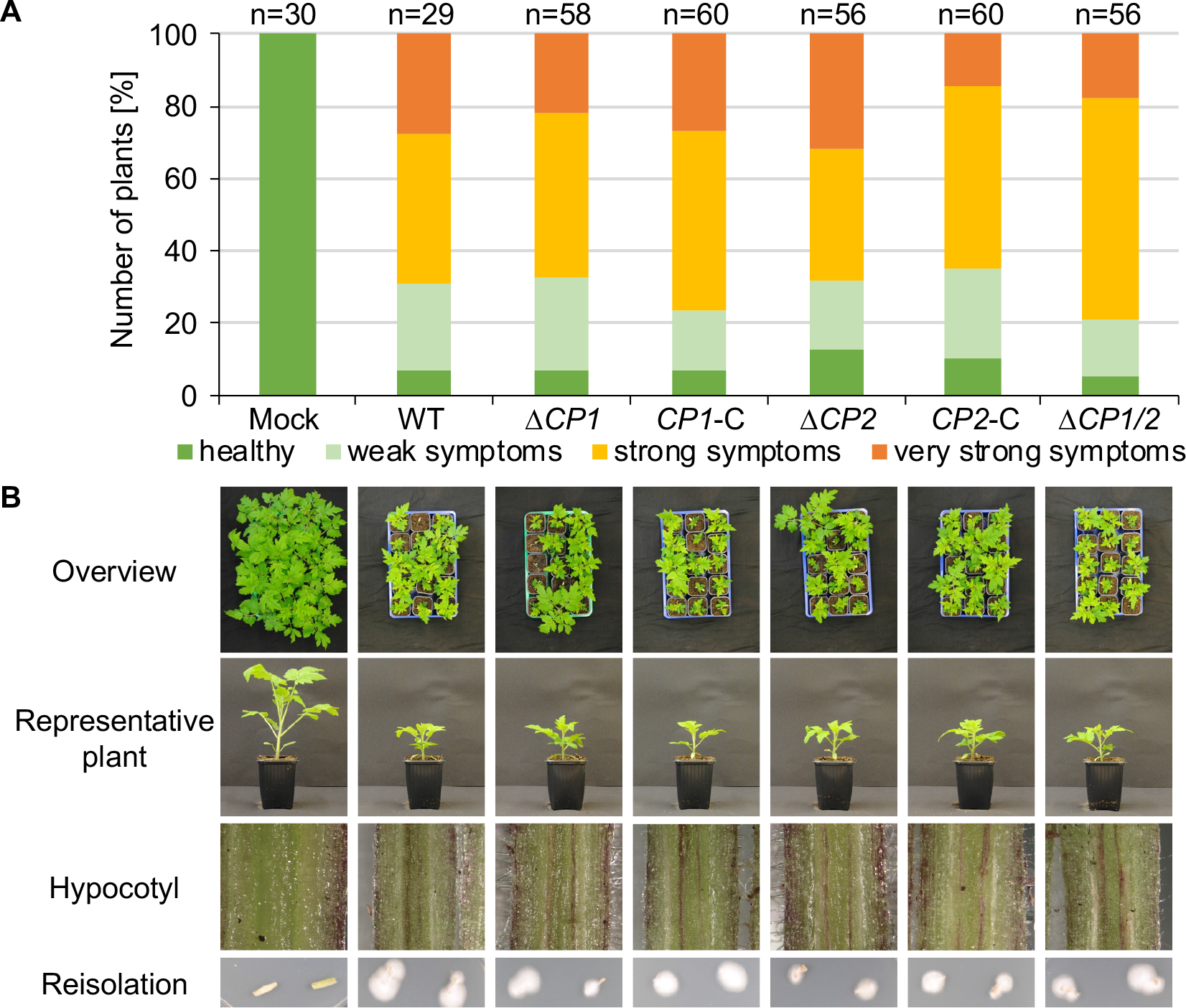
*V. dahliae* Cp1 and Cp2 are dispensable for virulence on tomato. Ten-day-old tomato seedlings were infected by root-dipping in a spore suspension of *V. dahliae* JR2 (WT) and the indicated deletion (Δ*CP1*, ΔCP2, Δ*CP1/2*) and complementation strains (*CP1*-C, *CP2*-C). Plants were incubated in the climate chamber under 16h:8h light:dark at 22-25°C. **(A)** The stack diagram shows the percentage of plants with the respective disease index. The disease index was assessed at 21 days post inoculation and includes the plant height, the longest leaf length and the fresh weight, which was compared to uninfected (mock) plants. The number (n) of treated plants is given for each fungal strain. **(B)** Representative plants, discolorations of the hypocotyls, and the fungal outgrowth of the stems are shown. The experiment was repeated twice. *CP1* and *CP2* deletion strains infect tomato plants to the same extent as WT and the complementation strains.

These experiments demonstrated that the tested CAZys or two genes encoding metalloproteases or cerato-platanin proteins do not affect *V. dahliae* pathogenicity under the tested conditions.

### Nlp3-GFP is secreted into the extracellular space

As described above, two necrosis and ethylene inducing-like proteins were detected in the xylem sap-specific exoproteome. Members of this group are known to contribute to *V. dahliae* pathogenicity and to exhibit host-specific roles [26, 27, 41]. Nlp1 and Nlp2 contribute to *V. dahliae* JR2 virulence on tomato and *Arabidopsis* [26], whereas corresponding proteins in *V. dahliae* V592 did not alter virulence on cotton [27]. Additionally, Nlp3 did not show any cytotoxic activity on *N. benthamiana* and has not been further characterized [26]. Because our exoproteome approach identified Nlp2 and Nlp3 as specifically secreted in xylem sap, we subsequently carried out additional analyses of the roles of these proteins.

To monitor the secretion of Nlp3 in liquid media, the gene was fused with a C-terminal *GFP* tag to *NLP3* under the control of the constitutive strong *gpdA* promoter. *V. dahliae* JR2 and *NLP3* deletion strains ectopically overexpressing *NLP3-GFP* were confirmed by Southern hybridization (S1 Fig). Growth characteristics of the strains expressing *NLP3-GFP* were analyzed as described for the other deletion strains. Similar to the deletion strains, the *NLP3-GFP* overexpressing mutants did not show any significant growth variation in comparison to the WT strain under the tested conditions as shown on CDM plates (S5 Fig). Confocal microscopy of the *NLP3-GFP* strains confirmed the production of a GFP signal derived by the Nlp3-GFP fusion protein with an intracellular location at vacuoles (Fig 6A). The expression and secretion of the fusion protein was analyzed by western experiments using a 24 h-old SXM culture. Intracellular proteins were extracted from fungal mycelium, extracellular proteins were precipitated from the culture supernatant and subjected to SDS-PAGE. Western analysis of the *NLP3-GFP* strain confirmed the overexpression and secretion of the fusion protein in pectin-rich medium (Fig 6B). WT and *NLP3* deletion strains ectopically overexpressing *NLP3-GFP* (WT*/OE-NLP3-GFP* and Δ*NLP3/OE-NLP3-GFP,* respectively) revealed strong signals for Nlp3-GFP with a size of 52 kDa in extracellular extracts after probing with *α*-GFP antibody whereas intracellular protein extracts result in faint bands at the size of the fusion protein as well as for free GFP (∼27 kDa). For the control strain *V. dahliae* JR2 expressing free GFP (WT/OE-*GFP*) no signal was detected in extracellular space and a strong GFP signal was detected in intracellular extracts. These data corroborate that Nlp3 is primarily a secreted protein and its expression levels neither influence growth nor development.

**Fig 6.**
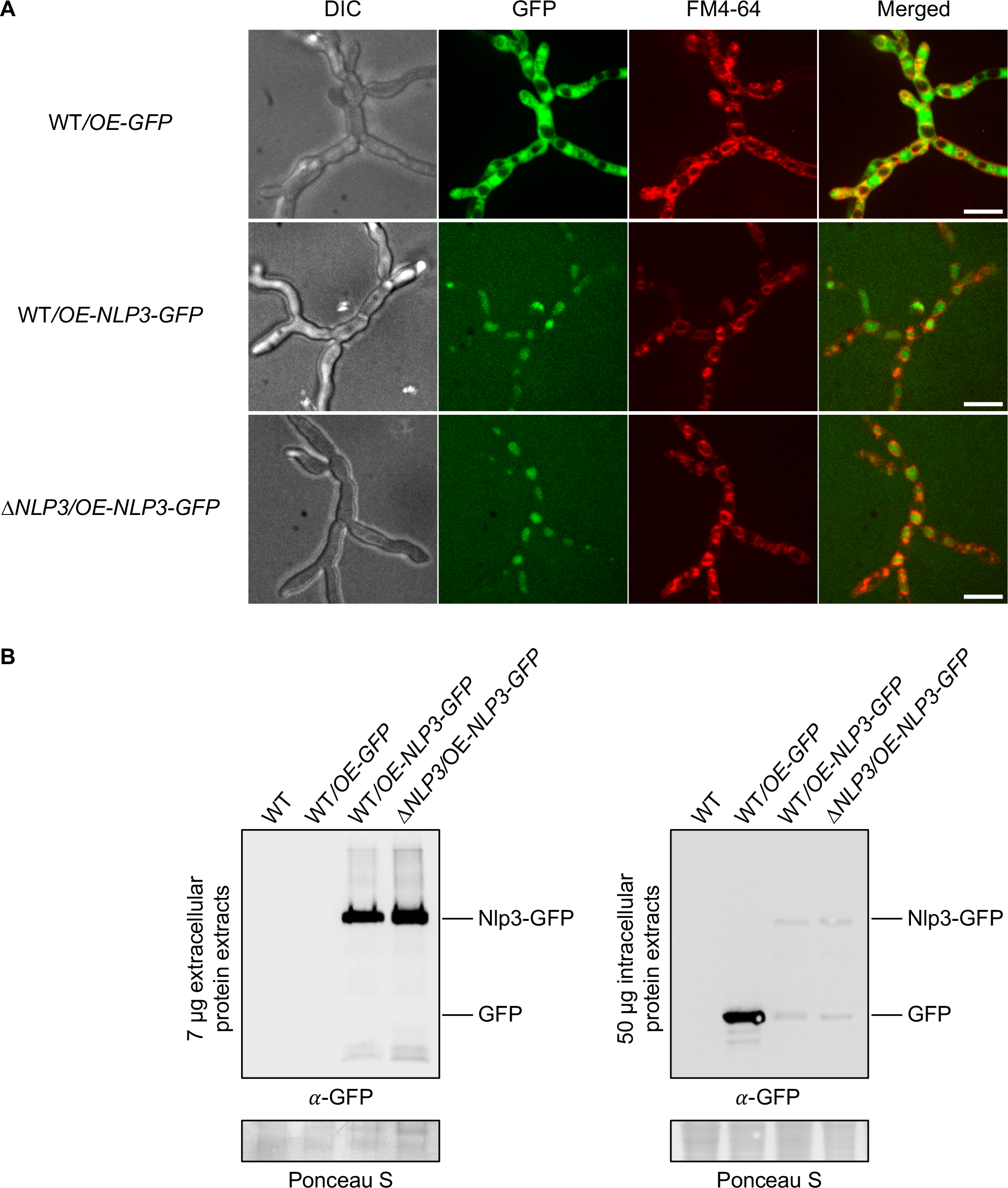
Nlp3-GFP is secreted into the extracellular space. *V. dahliae* JR2 (WT) ectopically overexpressing *GFP* (WT/OE-*GFP*) and WT and *NLP3* deletion strains ectopically overexpressing *NLP3-GFP* (WT/OE-*NLP3-GFP* and Δ*NLP3*/OE-*NLP3-GFP*, respectively) were tested for subcellular localization by fluorescence microscopy in **A** and localization and production of the intact full-length fusion protein intra- and extracellular by western hybridization in **B**. **(A)** Confocal microscopy of WT/OE-*NLP3-GFP* and Δ*NLP3*/OE-*NLP3-GFP* strains show the accumulation of the GFP signal inside the red-stained vacuoles whereas the strain WT/OE-*GFP* exhibits GFP signals in the cytoplasm. Spores of the indicated fungal strains were inoculated in 300 µl liquid PDM in µ-slide 8 well microcopy chambers (Ibidi) and incubated at 25°C overnight. Fungal hyphae were stained with the membrane-selective styryl dye *N*-(3-triethylammoniumpropyl)-4-(*p*-diethylaminophenyl-hexatrienyl) pyridinium dibromide (FM4-64). Scale bar = 10 µm. **(B)** Fungal strains grown in liquid simulated xylem medium (SXM) at 25°C for 24 hours. Western hybridization with *α*-GFP antibody was performed with 7 µg extracellular protein extracts from the culture supernatant and 50 µg intracellular protein extracts. Ponceau S staining served as loading control and WT was used as negative control. Analysis of extracellular proteins in the supernatant of WT/OE-*NLP3-GFP* and Δ*NLP3*/OE-*NLP3-GFP* strains revealed strong signals for Nlp3-GFP with a size of 52 kDa whereas intracellular protein extracts result in faint bands at the size of the fusion protein as well as for free GFP (∼27 kDa). In the control strain WT/OE-*GFP* a strong GFP signal was detected in intracellular extracts.

### Necrosis and ethylene inducing-like proteins Nlp2 and Nlp3 contribute to *V. dahliae* virulence on tomato

Nlp2 only exhibited minor effects in the *V. dahliae*-tomato system [26] whereas Nlp3 has not been tested for *V. dahliae* pathogenicity. We first analyzed the adherence to the root and further root colonization of the *NLP3* deletion strain on *A. thaliana* with fluorescence microscopy. The *NLP3* deletion strain expressing free *GFP* under the control of the *gpdA* promoter (Δ*NLP3/OE-GFP*) and the WT control overexpressing *GFP* (WT*/OE-GFP*) were used for root inoculation of three-week-old *Arabidopsis* seedlings. The root colonization at three and five days post inoculation was indistinguishable between WT*/OE-GFP* and Δ*NLP3/OE-GFP* (Fig 7A). Initial root colonization was observed at three days following inoculation and whole roots were covered with fungal hyphae after five days suggesting that Nlp3 is dispensable for *A. thaliana* root colonization.

**Fig 7.**
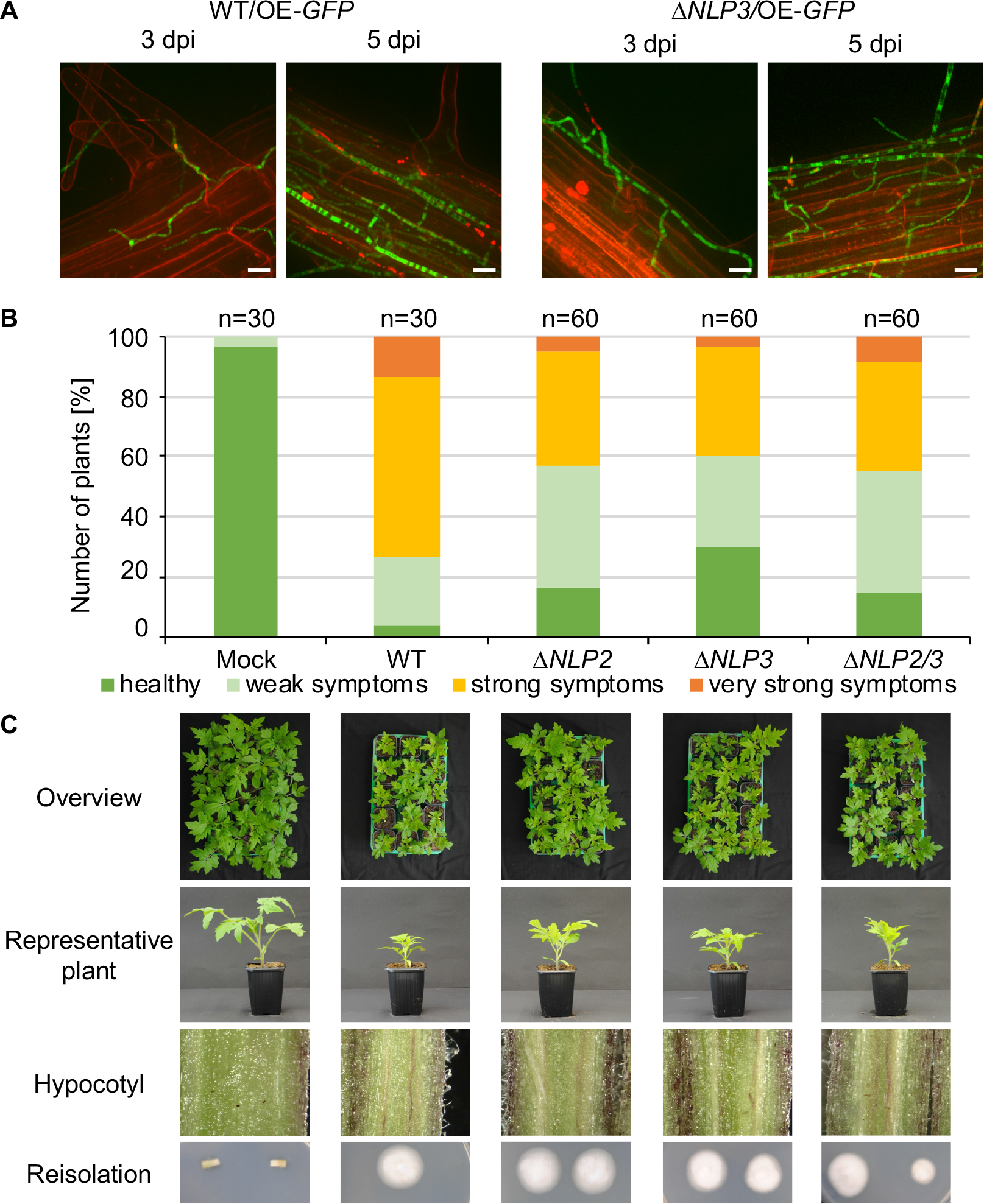
Necrosis and ethylene inducing-like effectors Nlp2 and Nlp3 contribute to *V. dahliae* virulence on tomato. The *NLP3* deletion strain was tested for *A. thaliana* root colonization and single and double deletion strains of *NLP2* and *NLP3* were tested for pathogenicity on tomato. Infected plants were incubated in the climate chamber under 16h:8h light:dark at 22-25°C. **(A)** Three-week-old *A. thaliana* seedlings were root-infected with spores of *V. dahliae* JR2 and *NLP3* deletion strain overexpressing free GFP (WT/OE-*GFP* and Δ*NLP3/*OE-*GFP*, respectively). At three and five days post inoculation (dpi) the colonization of the fungal hyphae was monitored with root cells stained by 0.05% propidium iodide/0.01% silwet solution. The experiment was repeated twice with two individual transformants of Δ*NLP3/*OE-*GFP*. Fluorescence microscopy pictures show similar initial colonization of *V. dahliae* WT/OE-*GFP* and Δ*NLP3/*OE-*GFP* on the root surface at 3 dpi and whole root colonization at 5 dpi. 3D Surface views were generated by Slidebook 5.0 software from stacks of single pictures. Scale bar = 10 µm. **(B, C)** Ten-day-old tomato seedlings were root-infected with spores of *V. dahliae* JR2 and the indicated single *NLP2* (Δ*NLP2*) and *NLP3* (Δ*NLP3*) and *NLP2/3* double deletion strains (Δ*NLP2/3*). Uninfected plants (mock) served as control. Representative plants, discolorations of the hypocotyls, and the fungal outgrowth of the surface sterilized stems are shown. The disease index was assessed at 21 dpi, which shows that plants infected with *NLP2* and *NLP3* single and double deletion strains exhibit an intermediate phenotype when compared with mock and WT-infected plants. The number of treated plants (n) is shown for each fungal strain. The experiment was repeated twice. For the deletion strains two individual transformants were tested.

Furthermore, the effect on pathogenicity towards tomato was investigated. Tomato infections were carried out as described above. All tested deletion strains were compromised in virulence compared to WT, but the plants nevertheless developed disease symptoms (Fig 7B). All infected plants exhibited stem discolorations and fungal outgrowth was detected from surface sterilized stems (Fig 7C, bottom row). Symptom development in plants colonized with deletion strains was less severe compared to WT infection. An overview of the trays with 15 treated plants, which is a representative number of plants considering fluctuations in the infection success, nicely demonstrate the differences between different strains (Fig 7C, top row). A representative plant also demonstrated the less stunted phenotype of plants treated with *NLP2* and *NLP3* single and double deletion strains in comparison to WT infected plants (Fig 7C, 2nd row). Translating the disease symptoms into the different categories revealed that about 60% of tested plants exhibited no or only mild symptoms compared to approximately 25% of the WT-treated plants (Fig 7B). Infections with the Δ*NLP2ΔNLP3* strain resulted in a similar disease index compared to the single deletion strains and, therefore, showed no additive effect of the two deleted *NLP* genes.

In conclusion, the infection study demonstrated that *NLP2* and *NLP3* contribute to *V. dahliae* JR2 virulence on tomato. Deletion of the genes still resulted in induction of disease symptoms suggesting that the fungal strains are well able to penetrate the plant and that Nlp2 and Nlp3 play primarily a role inside of the plant. This is consistent with the exoproteome approach that identified Nlp2 and Nlp3 as xylem sap-specific secretion proteins, in accordance with a role during later infection steps in the xylem vessels. These experiments show that our proteomic approach successfully identified a xylem sap-specific group that includes proteins that are uniquely required in the xylem sap of the plant. While other tested proteins may have redundant functions, we were able to identify NLPs, which are only secreted in a specific environment, as candidates important for *Verticillium* infection.

## Discussion

Fungi require sensing and adapting mechanisms throughout their life cycle. Different environmental cues induce different secretion responses enabling the pathogen to react to changes in e.g. nutrient supply or host defense responses [42].

Our experiments provide evidence for the ability of the allodiploid *V. longisporum* to distinguish between different environments and to induce media-dependent secretion responses. *V. longisporum* secretes a general protein response pattern in various non plant-related media, which reflects a situation outside of the plant. During cultivation in pectin-rich SXM or plant-extracted xylem sap, the fungus reacts to its surrounding and secretes specific proteins important for the degradation of plant material and the colonization of the xylem. These results imply a complex recognition of plant material in the environment and further show that SXM lacks the full capacity to mimic the natural growth medium xylem sap.

Xylem sap consists of water, plant defense proteins, hormones and low concentrations of amino acids and sugars that are transported to upper parts of the plant through the transpiration stream [7, 8]. SXM was developed to mimic the *in planta* environment, but mainly contains amino acids and the complex carbon source pectin [32]. Pectin is found in plant cell walls where it strengthens the wall integrity [43]. The degradation of this complex branched polysaccharide demands the action of several carbohydrate-active enzymes [39]. This situation is reflected in our SXM-derived exoproteome. In the SXM-specific and the core exoproteome we found 64 and 90 CAZys, respectively, of which the enzymes acting on pectin are especially overrepresented with 32 and 20 proteins. Other studies examined the upregulation of fungal genes after *V. dahliae* root-inoculation of *A. thaliana* seedlings for one day [44]. Of these upregulated genes, we identified corresponding proteins are found in our xylem sap-specific as well as the SXM-specific and core exoproteome. Another study on the *V. dahliae* secretome used minimal medium with cotton root fragments [22]. Several secreted proteins were identified including 12 cellulases, five pectate lyases, two chitinases, 13 proteases and one cerato-platanin domain containing protein (Cp1) [22]. A number of these proteins were detected in our SXM-specific and core exoproteome. However, no overlap to our xylem sap-specific secretome was detected suggesting that the SXM-induced response of *Verticillium* spp. is more similar to the presence of root fragments. In contrast, the exoproteome after contact with the living plant shows an overlap to our SXM- and xylem sap-derived exoproteomes.

The xylem sap is a unique niche for fungal growth due to its low and imbalanced nutrient supply [7, 8]. It therefore is likely quite important for *Verticillium* to recognize this specific environment and adapt to it by secreting colonization-related proteins. In prior studies, we demonstrated that *V. longisporum* changes its growth according to the presence or absence of host xylem sap [31]. The fungus is able to adapt to the low-nutrient and imbalanced amino acid supply in the xylem sap by activating the cross-pathway [10]. In filamentous fungi, this process is controlled by the cross-pathway control transcription factor Cpc1, which is encoded by a homolog of the yeast gene *GCN4* (general control non-derepressed) [10, 45, 46]. Knockdowns in *V. longisporum* and knockouts in *V. dahliae* revealed that Cpc1 is required for growth under amino acid starvation conditions and successful colonization of the host plants [10]. These findings show that *V. longisporum* senses and reacts to its host environment to survive.

The haploid *V. dahliae* responds differently in a susceptible and tolerant olive cultivar [47]. In the susceptible cultivar, the fungus significantly induced expression of genes involved in niche-adaptation, pathogenicity and microsclerotia development [47]. Similarly, the transcriptome of two *V. dahliae* strains with different virulence levels on cotton were analyzed. The strain with reduced virulence exhibited more repressed genes, of which most are related to pathogenesis [48]. These results corroborate that the fungus senses its environment and responds with different secretion patterns.

Another tight and fast adapting control mechanism of gene expression lies in chromatin modifications, which can be induced by environmental changes [49]. Such an epigenetic-mediated control has been observed for effector expression in *Leptosphaeria maculans.* Effector genes often reside in AT-rich regions of the genome. These are associated with heterochromatin and explain the silenced state of effector expression. Upon leaf infection chromatin-mediated repression is abolished and gene expression is upregulated in *L. maculans* [50, 51]. As *V. longisporum* responds to the presence of plant-related compounds by inducing specific exoproteome patterns, it will be interesting to shed light on a putative chromatin modification contribution to these unique responses.

Transcriptional regulators can induce the gene expression of several effectors at once. For example, the transcription factors Som1, Vta2 and Vta3, that are required for sequential steps of infection, control similar but also distinct sets of secreted proteins involved in virulence [52, 53]. All three proteins are involved in the regulation of, for example, *NLP2* [52, 53]. The expression of the two cytotoxic NLPs, *NLP1* and *NLP2*, from *V. dahliae* has been analyzed during host colonization. When colonizing tomato plants, both transcript levels were elevated although only *NLP1* expression was increased during colonization of tobacco plants. The *in planta* expression of *NLP1* and *NLP2* corresponds to the infection phenotype of the deletion strains [26]. These results confirm the hypothesis of a sensitive control mechanism. It shows that effectors may act host-specifically and are only expressed in suitable hosts supporting the idea of a fine-tuned response.

In this study, we identified two NLPs, Nlp2 and Nlp3, as effectors specifically secreted in xylem sap. Single and double deletions of the corresponding genes resulted in compromised pathogenicity on tomato. The *NLP2* single deletion strain was included as control as it was shown previously to contribute to *V. dahliae* virulence [26]. The *NLP3* deletion strain was additionally tested for *A. thaliana* root colonization revealing no differences to WT (Fig 7A). This indicates that Nlp3 is dispensable for root adhesion or colonization but is required during later steps of the infection. NLPs characteristically induce necrotic lesions [27] thus supporting their activity inside of the plant.

Other xylem sap-specifically enriched candidates did not show an effect on *V. dahliae* pathogenicity on tomato. This may be due to redundant or greater contributions of the hundreds of secreted *Verticillium* effectors [29]. Our approach revealed a high number of CAZys among the *V. longisporum* secretome representing 173 out of the total 399 identified proteins (S4 Table). Overall, the *V. dahliae* genome exhibits a strikingly high repertoire of CAZys, especially pectin-degrading enzymes [29]. This suggests that proteins with redundant functions are secreted and explains the WT-like infections of *V. dahliae* strains lacking one of the tested CAZys (glucoamylase Gla1, putative polysaccharide mono-oxygenase Cbd1 and *α*-amylase Amy1).

We identified two metalloproteases, Mep1 and Mep2, belonging to two different groups of metalloproteases (M36 and M43, respectively) as xylem sap-specific secreted proteins. Metalloproteases promote fungal virulence by degrading host proteins [54, 55]. Plant chitinases degrade chitin of the fungus, which elicits the plant defense response [56]. *V. dahliae* possesses the ability to truncate extracellular chitin-binding domain-containing chitinases [24, 54]. The *V. dahliae* proteome comprises two M43 and six M36 peptidases implicating that other metallopeptidases are able to complement the absence of Mep1 and Mep2. Furthermore, synergistic actions of metallo- and serine proteases have been reported in *F. oxysporum* [54], providing more evidence for the hypothesis of functional redundancy.

We also investigated on the cerato-platanin domain containing proteins Cp1 and Cp2. Cp1 was first identified in the exoproteome of the *V. dahliae* strain XH-8 when incubated in minimal medium supplemented with cotton root fragments. In this system, Cp1 was required for cotton virulence and is suggested to function as a chitin scavenger to prevent fungal recognition by the plant [22]. We detected Cp1 in our core exoproteome and included it in our study to investigate on putative synergistic actions of the two CPs present in the *V. dahliae* genome. Our results show that Cp1 and Cp2 are dispensable for *V. dahliae* pathogenicity on tomato. These results indicate that effectors may have strain- and host-specific activities.

To our knowledge, our proteomic study is the first report identifying the differences between the secretion responses of *V. longisporum* in host xylem sap, the xylem sap mimic SXM and other media. Non-plant related environments elicited a similar broad exoproteome pattern whereas the plant-related media, SXM and xylem sap, induced similar but also distinct responses. These results indicated that the fungus has the capacity to sense differences in the presence of plant-related compounds and therefore rules out SXM as xylem sap mimic. Additionally, our approach identified necrosis and ethylene inducing-like proteins Nlp2 and Nlp3 in the xylem sap-specific secretome. These proteins are required for *V. dahliae* pathogenicity with roles in later steps of infection and display potential targets for control strategies of Verticillium wilt.

## Materials and Methods

### Fungal strains and growth conditions

*Verticillium* strains (S5 Table) were cultivated in liquid simulated xylem medium (SXM) modified from [32] as described in [57] for conidiospore formation and in liquid potato dextrose medium (PDM) (Potato Dextrose broth (Carl Roth)) for mycelial growth. Cultures were incubated at 25°C under constant agitation at 115 – 125 rpm. For long-term storage spores were maintained in closed vials with 25% glycerol at −80°C. For the exoproteome comparison, *V. longisporum* 43 (Vl43) was inoculated with 1.5 x 10^6^ spores per 150 ml PDM (150 rpm) and incubated for four days. Each culture was centrifuged and the mycelium and spore sediment was resuspended in 150 ml extracted xylem sap of *B. napus*; SXM, the minimal medium Czapek-Dox medium (CDM, modified from [58] and [59]) supplemented with 7% extracted xylem sap or plant proteins; H_2_O and H_2_O supplemented with 0.1% glucose; YNB (yeast nitrogen base: 1.5 g/l YNB, 5 g/l (NH_4_)2SO_4_, 20 g/l glucose, ad 1 l H_2_O) and vegetable juice (V8, Campbell Soup Company). After an additional incubation period of four days, proteins of the supernatant were precipitated with TCA/acetone.

### Xylem sap extraction

Xylem sap was extracted from *B. napus* (Falcon, Norddeutsche Pflanzensucht). Seeds were surface sterilized with 70% ethanol and sown on sand. Plants were grown at long-day condition (16h light: 8h dark) and 22°C. Seven-day-old seedlings were transferred into a soil/sand (1: 1) mixture and grown for 42 days. To extract the xylem sap, plants were cut at the height of the first internode and the xylem sap was collected. The xylem sap was filtered through Vivaspin 15R centrifugal concentrators (Sartorius) and directly used as medium for inoculation.

### Genomic DNA Extraction

For isolation of genomic DNA (gDNA) from fungal powder, the method modified from [60] was used. The fine powder was mixed with 800 µl of lysis buffer (50 mM Tris (pH 7.5), 50 mM EDTA (pH 8), 3% (w/v) SDS and 1% (v/v) ß-mercaptoethanol) and incubated at 65°C for one hour. Before the mixture was centrifuged for 20 min at 13000 rpm, 800 µl phenol were added. The upper aqueous phase was transferred to a new tube. To denature the proteins, 500 μl chloroform were added, mixed and centrifuged for 10 min at 13000 rpm. The upper phase was mixed with 400 µl of isopropanol for precipitation of gDNA and centrifuged for 2 min at 13000 rpm. The sedimented gDNA was washed with 70% (v/v) ethanol. The gDNA was dried at 65°C for approximately 25 min before it was dissolved in up to 100 µl deionized H_2_O containing 2 µl RNase A (10 mg/ml) and treated at 65°C for 30 min to remove RNA.

### Plasmid and strain construction

The desired genes and flanking regions for plasmid construction were amplified of *V. dahliae* JR2 WT gDNA with the Phusion High Fidelity Polymerase, *Taq* DNA Polymerase (both Thermo Fisher Scientific) or Q5 High Fidelity Polymerase (New England Biolabs). Primers are listed in S6 Table.

GeneArt Seamless Cloning and Assembly Kit (Thermo Fisher Scientific) was used for the cloning strategy. Plasmids are listed in S7 Table. *E. coli* strain DH5*α* was utilized for cloning reactions and propagation of plasmids. Transformation of *E. coli* was performed based on a heat shock method [61]. *A. tumefaciens* AGL-1 cells were transformed with the desired plasmids via a freeze-thaw method [62] and was then utilized for an *A. tumefaciens* mediated transformation of *V. dahliae* spores, which was performed based on the method described by [63]. Details on specific mutant strains in haploid *V. dahliae* are given in S1 Text.

### Southern hybridization analysis

For verification of *V. dahliae* deletion strains, the corresponding flanking region of the gene was amplified and labeled as probe. Genomic DNAs were restricted with indicated enzymes overnight. The mixture was separated on a 1% agarose gel, and DNA was transferred to a Hybond-N membrane (GE Healthcare) by blotting. DNA on the membrane was hybridized overnight to the probe. CDP-Star Detection reagent (GE Healthcare) was used to detect chemiluminescence signals according to the manufacturer’s instructions.

### Protein assays and western hybridization analysis

Extracellular proteins from the supernatant of SXM cultures were precipitated with 10% TCA (w/v) in acetone at 4°C overnight. This mixture was centrifuged at 4 000 rpm for 60 min at 4°C. The protein sediment was washed three times with 80% (v/v) acetone, once with 100% (v/v) acetone and then dissolved in 8 M urea/ 2 M thiourea. Intracellular proteins were extracted from ground mycelium with extraction buffer (300 mM NaCl, 100 mM Tris-HCl pH 7.5, 10% glycerol, 1 mM EDTA, 0.02% NP-40, 2 mM DTT and complete EDTA-free protease inhibitor cocktail (Roche)). Samples were centrifuged for 20 min at 13 000 rpm at 4°C and the supernatants were transferred into fresh test tubes. Protein concentrations were determined using a Bradford-based Roti-Quant assay (Carl Roth). Protein samples were separated in 12% SDS-PAGE gels, followed by protein transfer onto an Amersham Protran 0.45 µm nitrocellulose membrane (GE Healthcare). The membrane was blocked in 5% (w/v) skim milk powder in TBS-T (10 mM Tris-HCl (pH 8), 150 mM NaCl, 0.05% (w/v) Tween 20) and probed with α-GFP antibody (Santa Cruz Biotechnology). As secondary antibody the horseradish peroxidase-coupled α-mouse antibody (115-035-003, Jackson ImmunoResearch) was applied. Detection of chemiluminescent signals was conducted with horseradish peroxidase substrate luminol based chemiluminescence.

### Tryptic digestion, mass spectrometry analysis and protein identification

For one-dimensional gel analysis 30 µg of the extracellular protein extract was separated by 12% SDS-PAGE gels. The polyacrylamide gels were incubated 1h in fixing solution (40% (v/v) ethanol, 10% (v/v) acetic acid) and washed twice for 20 min with H_2_O. Gels were colloidal Coomassie stained (0.12% (w/v) CBB G-250, 5% (w/v) aluminum sulphate-(14-18)-hydrate, 10% (v/v) methanol, 2% (v/v) orthophosphoric acid (85%)) and two lanes for each growth condition were cut into ten pieces of equal size. The excised polyacrylamide gels were in-gel digested with trypsin [64]. Resulting tryptic peptide mixtures were separated by a reversed-phase liquid chromatographic column (Acclaim PepMap RSLC column, 75 μm x 15 cm, C18, 3 μm, 100 Å, P/N164534, Thermo Fisher Scientific) to further reduce sample complexity prior to mass analyses with an LTQ Velos Pro mass spectrometer (Thermo Fisher Scientific). To identify matches of the detected peptides an in-house genome-wide protein sequence database of *Verticillium longisporum* was used of which the protein sequences are given in S1 Table. Analysis was performed by a Thermo Proteome Discoverer (version 1.3) workflow that integrates Sequest and Mascot search engines. For the search an initial precursor mass tolerance of 10 ppm and fragment mass tolerance of 0.8 Da Carbamidomethylcysteine was used as fixed modification. Oxidized methionine was included as variable modification and two miscleavages were allowed for each peptide. For peptide and protein validation, a 0.5% false discovery rate was set and determined by using peptide validator with a reverse decoy database. Resulting lists of identified proteins were semi-quantitatively processed using the Marvis Suite software [34]. Only proteins with a WoLF PSORT extracellular localization prediction of above 12 were considered for further characterization [65]. A group of 445 identified proteins fulfilled the criteria of a threshold of 1 peptide and a high intensity ratio of 0.83 in one condition. The protein sequences of the 445 proteins, and details on their abundance as measured by identified peptides, are given in S1 Table. The signal-to-level (s/l) ratio (see MarVis-Suite handbook on http://marvis.gobics.de) was calculated using as signal the difference between SXM and xylem sap condition averages (or vice versa) and as level the corresponding maximum. Polypeptides with an s/l ratio above 0.3 were considered as candidates with higher intensities in the specific medium and therefore belong to the specifically enriched proteins. Xylem sap-specific proteins are depicted in green and SXM-specific proteins in red. Whereas an s/l ratio below 0.3 was considered not to be specifically enriched and formed the shared core proteome.

The list of proteins was further analyzed and compared to the Ensembl Fungi annotations which are more robust. Candidates with no proper predictions in our preliminary genome-wide protein sequence database of *V. longisporum* (e.g. stop codons in protein sequences, two genes annotated as one) were revealed by comparison with corresponding *V. dahliae* JR2 and *V. alfalfae* VaMs.102 sequences and eliminated from the list. Further analyses are based on the *V. dahliae* JR2 or the *V. alfalfae* VaMs.102 protein sequences [35]. Domain predictions and associated families were obtained with InterProScan [36] and classification of carbohydrate-active enzyme (CAZys) families according to the CAZy database (http://www.cazy.org) was specified with dbCAN2 [37] to address putative functions of the proteins. Proteins were considered as putatively secreted with at least one predicted signal peptide by InterProScan [36] or SignalP-5.0 [66] or as long as it passed the threshold of 12 as determined by WoLF PSORT [65] for the *V. longisporum* 43, *V. dahliae* JR2 or *V. alfalfae* VaMs.102 protein sequence. All details are given in S2 Table.

### *A. thaliana* root infection assay

*A. thaliana* (Col-0, N1902; Nottingham Arabidopsis Stock Centre) seedlings were inoculated by root-dipping with a conidia suspension of *V. dahliae* JR2 and *NLP3* deletion strain overexpressing *GFP* (1×10^7^ spores/ml) based on the method described by colleagues [53]. Three-week-old seedlings were used for infection. The roots were incubated in spore solutions with 100 000 spores/ml for 35 minutes. The plates were further incubated in the plant chamber at long day conditions (22-25°C) and colonization on the roots was monitored at indicated time points. The root was incubated in a staining solution (0.0025% (v/v) propidium iodide, 0.005% (v/v) silwet) for five minutes in the dark. Images of infected roots were taken with a Zeiss Observer Z1 microscope equipped with CSU-X1 A1 confocal scanner unit (Yokogawa), QuandtEM:512SC (Photometrics) digital camera and Slidebook 5.0 software package (Intelligent Imaging Innovations).

### Tomato infection assay

*Solanum lycopersicum* (Moneymaker, Bruno Nebelung Kiepenkerl-Pflanzenzüchtung) seeds were surface sterilized with 70% (v/v) EtOH, 0.05% Tween 20, sown on sand/soil (1:1) mixture (Dorsilit, Archut). Plants were grown under a photoperiod of 16 h light and 8 h of darkness at 25 and 22°C, respectively. The tomato pathogenicity assays were performed on ten-day-old *S. lycopersicum* seedlings. The plants were root-inoculated by incubating the roots in 50 ml of 10^7^ spores/ml for 40 min under constant agitation at ∼35 rpm. Mock control plants were treated similarly with dH_2_O.

The seedlings were transferred to pots containing a sand/soil (1:1) mixture and 3 000 000 spores or 3 ml dH_2_O for mock plants were added to the soil. For each strain or control 15 plants were infected. Plants were incubated in the climate chamber at long day conditions for another 21 days before disease symptoms were scored. The fresh weight of the aerial parts, the length of the longest leaf and the height of the vegetation point were measured. These parameters were calculated into a disease score ranking relative to uninfected mock plants. The mean values of mock plants of each parameter were set to 100%. All values above 80% were assigned as ‘healthy’, 60-80% as ‘mild symptoms’, 40-80% as ‘strong symptoms’, lower than 40% as ‘very strong symptoms’ and dead plants as ‘dead’. The scores for each strain were visualized in a stack diagram displaying the number of plants per disease score relative to the total amount of tested plants from all experiments. As another measure the discoloration of the tomato hypocotyl was observed with a binocular microscope SZX12-ILLB2-200 from Olympus. All treated plants were tested for fungal outgrowth 21 days after infection. The tomato stems were surface sterilized in 70% ethanol, followed by 6% hypochlorite solution each for 8 min before two washing steps with dH_2_O. Stem ends were removed and slices were placed on PDM plates containing 100 µg/ml chloramphenicol. After incubation of seven days at 25°C the fungal outgrowth was observed. The pathogenicity assay was performed once with the metalloprotease constructs and once with Δ*GLA1*, Δ*CBD1* and Δ*AMY1* strains. Pathogenicity assays with the *CP* and *NLP* constructs were repeated twice. For each assay 15 plants were infected with WT and 15 plants were used as mock control. Per transformant 15 plants were infected and scores of identical strains were taken together of two individual transformants.

### Accession numbers

Sequence data for *V. dahliae* were retrieved from Ensembl Fungi with the following accession numbers: *NLP2* (*VDAG_JR2_Chr2g05460a*), *NLP3* (*VDAG_JR2_Chr4g05950a*), *CP1* (*VDAG_JR2_Chr7g00860a*), *CP2* (*VDAG_JR2_Chr2g07000a*)*, MEP1* (VDAG_JR2_Chr8g09760a), *MEP2* (VDAG_JR2_Chr1g21900a), *GLA1* (*VDAG_JR2_Chr8g11020a*), *CBD1* (*VDAG_JR2_Chr4g04440a*), *AMY1* (*VDAG_JR2_Chr7g03330a*). Protein sequences for *V. longisporum* are given in S1 Table as retrieved from an in-house database or taken from Ensembl Fungi.

## Acknowledgments

The authors thank N. Scheiter for technical assistance and A. Keyl and A. Pelizaeus for help with the experimental work during lab rotations. This research has been funded by the German Research Foundation (DFG, IRTG 2172 “PRoTECT” program of the Göttingen Graduate Center of Neurosciences, Biophysics, and Molecular Biosciences) to ML, JS, IM, AN, GHB, and IF, the Federal Ministry of Education and Research (BMBF “BioFung”, FKZ 0315595) to AK, ML, SB-S, GHB, BM and IF and an NSERC CREATE award to JWK.

## Supporting information

**S1 Fig. Verification of *V. dahliae NLP2* and *NLP3* deletion or overexpression constructs.** *V. dahliae* JR2 (WT) was transformed with the respective deletion cassettes to generate *NLP2* and *NLP3* deletion strains (Δ*NLP2* and Δ*NLP3*, respectively) via homologous recombination. The *NLP2/NLP3* double deletion strain (Δ*NLP2/3*) was obtained by transforming the Δ*NLP3* strain with the Δ*NLP2* construct. Strains overexpressing ectopically integrated *GFP* or *NLP3-GFP* were constructed by transforming WT or *NLP3* deletion strains with the displayed constructs (*GFP* and *NLP3-GFP* are driven by the *gpdA* promoter and followed by the *trpC* terminator). All constructs contain resistance cassettes (*HYG^R^*: hygromycin resistance; *NAT^R^*: nourseothricin resistance) with a *gpdA* promoter and a *trpC* terminator. Schemes with used restriction enzymes, corresponding cutting sites (arrows) and probes (red lines) used for Southern hybridization are presented in **A** and **B**. **(A)** *XhoI* and the 5’ flanking region as probe were used for verification of the *NLP2* deletion strain. **(B)** Δ*NLP3* strain was confirmed using *Xho*I with 5’ flanking region as probe. And ectopic integration of *GFP* into Δ*NLP3* strain and *NLP3-GFP* into WT or Δ*NLP3* strain was verified with restriction enzyme *Bgl*I with *GFP* as probe. **(C)** Confirmation of deletion strains by Southern hybridization is shown for Δ*NLP2* #4 = VGB390, #8 = VGB391; Δ*NLP3* #4 = VGB384, #5 = VGB385, Δ*NLP2/3* #6 = VGB400, #7 = VGB401; Δ*NLP3/*OE*-GFP* #8 = VGB431, #9 = VGB432, WT/OE-*NLP3-GFP* #2 = VGB407, #6 = VGB408, Δ*NLP3/*OE*-NLP3-GFP* #1 = VGB409, #7 = VGB410. Genomic WT DNA served as control. Restriction enzymes, probes and sizes of expected fragments are indicated.

**S2 Fig. Verification of *V. dahliae CP1* and *CP2* deletion and complementation strains.** Deletion strains (Δ*CP1 and* Δ*CP2*) were obtained via homologous recombination between the deletion constructs and *V. dahliae* JR2 (WT). The *CP2* deletion strain was transformed with the *CP1* deletion construct to generate the double deletion strain Δ*CP1/2*. To generate ectopic complementation strains the constructs were integrated into the deletion strain as indicated by //. The constructs contain resistance cassettes (*HYG^R^*: hygromycin resistance; *NAT^R^*: nourseothricin resistance) controlled by the *gpdA* promoter and the *trpC* terminator. Schemes with used restriction enzymes, corresponding cutting sites (arrows) and probes (red lines) used for Southern hybridization are presented in **A** and **B**. **(A)** Confirmation of *CP1* deletion and complementation strain was achieved by enzyme restriction of gDNA with *Sal*I and 3’ flanking region as probe or *Mfe*I and 5’ flanking region as probe. **(B)** *Sma*I with 3’ flanking region or *Hind*III with 5’ flanking region as probe was used for *CP2* deletion and complementation strains. **(C)** Deletion and complementation strains were verified by Southern hybridization: Δ*CP1 #*11 = VGB316, #13 = VGB317; *CP1*-C #6 = VGB489, #7 = VGB490; Δ*CP2* #4 = VGB406, #14 = VGB422; *CP2*-C #3 = VGB429, #7 = VGB430; Δ*CP1*Δ*CP2* #5 = VGB423, #10 = VGB424. Genomic WT DNA was used as control. Restriction enzymes, probes and sizes of expected fragments are depicted.

**S3 Fig. Verification of *V. dahliae MEP1* and *MEP2* deletion strains.** *MEP1* and *MEP2* deletion strains (Δ*MEP1* and Δ*MEP2*, respectively) were constructed via homologous recombination between the deletion construct and *V. dahliae* JR2 (WT) and confirmed by Southern hybridization. Restriction sites of used enzymes (arrows) and probes (red lines) used for Southern hybridization are depicted in **A** and **B**. **(A)** Deletion of *MEP1* was confirmed after enzyme restriction of gDNA with *Kpn*I or *Bgl*II and 3’ flanking region as probe. *MEP1* deletion construct contains hygromycin resistance cassette (*HYG^R^*) controlled by the *gpdA* promoter and the *trpC* terminator. **(B)** Verification of *MEP2* deletion strain containing the nourseothricin resistance (*NAT^R^*) cassette under control of the *gpdA* promoter and the *trpC* terminator was achieved with *Sac*I or *Nco*I restriction and 3’ flanking region as probe. **(C)** Deletion strains were confirmed by Southern hybridization: Δ*MEP1* #7 = VGB226, #8 = VGB227; Δ*MEP2* #2 = VGB133, #8 = VGB126; Δ*MEP1/2* #2 = VGB204, #5 = VGB203. Genomic WT DNA served as control. Restriction enzymes, probes and sizes of expected fragments are indicated.

**S4 Fig. Verification of *V. dahliae GLA1*, *CBD1* and *AMY1* deletion strains.** By homologous recombination between the deletion construct and *V. dahliae* JR2 (WT) the depicted deletion strains were obtained. The deletion constructs contain a nourseothricin resistance cassette (*NAT^R^*) controlled by the *gpdA* promoter and the *trpC* terminator. The used probes (red lines) and restriction enzymes with their cutting sites (arrows) are illustrated in the schemes (left). The expected fragments were confirmed by Southern hybridization for each deletion strain with WT as control (right). **(A)** With *Xho*I and 5’ flanking region as probe *GLA1* deletion strains (Δ*GLA1*) #4 = VGB129 and #10 = VGB131 were verified. **(B)** Verification of *CBD1* deletion strains (Δ*CBD1* #2 = VGB127, #5 = VGB132) was achieved with *Pvu*I enzyme restriction and 3’ flanking region as probe and **(C)** *AMY1* deletion strains (Δ*AMY1* #2 = VGB130, #10 = VGB128) were verified with *Kpn*I and 3’ flanking region as probe.

**S5 Fig. Expression of *NLP3-GFP* under control of a constitutively active promoter allows *V. dahliae* wildtype-like growth on solid media.** 50 000 spores of *V. dahliae* JR2 wildtype (WT) and *NLP3* deletion strains (Δ*NLP3*) and corresponding strains ectopically overexpressing *NLP3-GFP* (WT/*OE-NLP3-GFP* and Δ*NLP3*/*OE-NLP3-GFP*, respectively) were point inoculated on minimal medium (CDM) plates. The growth phenotype was observed after 10 days of incubation at 25°C and revealed a similar phenotype for all tested strains.

S1 Table. List of identified proteins in SXM and xylem sap.

S2 Table. Annotation of identified proteins in SXM and xylem sap.

S3 Table. Functional groups of secreted proteins.

S4 Table. CAZy classification of secreted proteins.

S5 Table. *Verticillium* strains used in this study.

S6 Table. Primers used in this study.

S7 Table. Plasmids used in this study.

S1 Text. Additional Materials and Methods.

