## Supplementary Material for "*V. longisporum* elicits media-dependent secretome responses with a further capacity to distinguish between plant-related environments"

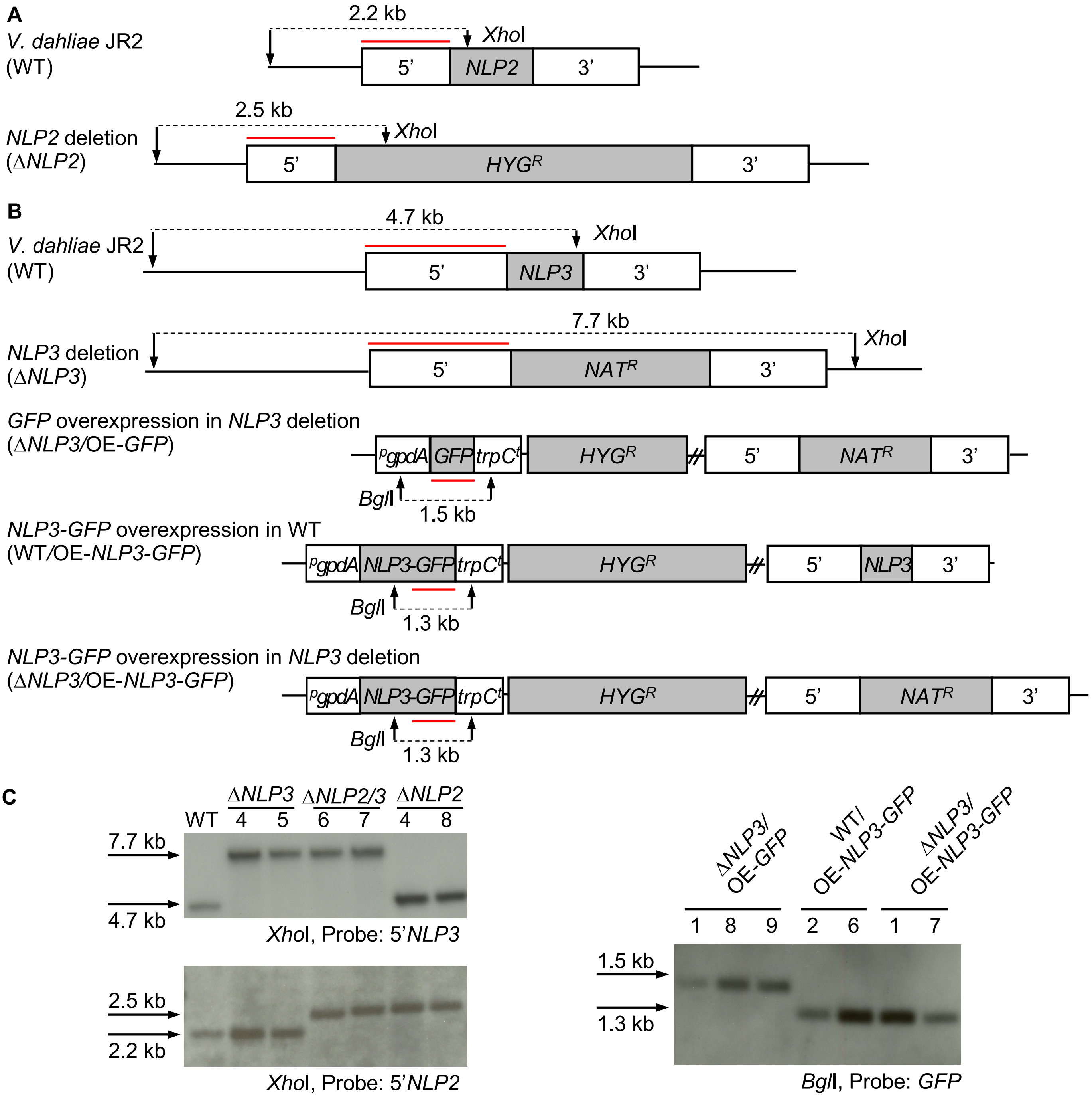
A
V. dahliae JR2 (WT)
2.2 kb
XhoI
5'
NLP2
3'
NLP2 deletion (ΔNLP2)
2.5 kb
XhoI
5'
HYGR
3'
B
V. dahliae JR2 (WT)
4.7 kb
XhoI
5'
NLP3
3'
NLP3 deletion (ΔNLP3)
7.7 kb
XhoI
5'
NATR
3'
GFP overexpression in NLP3 deletion (ΔNLP3/OE-GFP)
pgpdA
GFP
trpCt
HYGR
5'
NATR
3'
BglI
1.5 kb
NLP3-GFP overexpression in WT (WT/OE-NLP3-GFP)
pgpdA
NLP3-GFP
trpCt
HYGR
5'
NLP3
3'
BglI
1.3 kb
NLP3-GFP overexpression in NLP3 deletion (ΔNLP3/OE-NLP3-GFP)
pgpdA
NLP3-GFP
trpCt
HYGR
5'
NATR
3'
BglI
1.3 kb
C
WT
ΔNLP3 4 5
ΔNLP2/3 6 7
ΔNLP2 4 8
7.7 kb
4.7 kb
XhoI, Probe: 5'NLP3
2.5 kb
2.2 kb
XhoI, Probe: 5'NLP2
ΔNLP3/OE-GFP 1 8 9
WT/OE-NLP3-GFP 2 6
ΔNLP3/OE-NLP3-GFP 1 7
1.5 kb
1.3 kb
BglI, Probe: GFP

**S1 Figure**

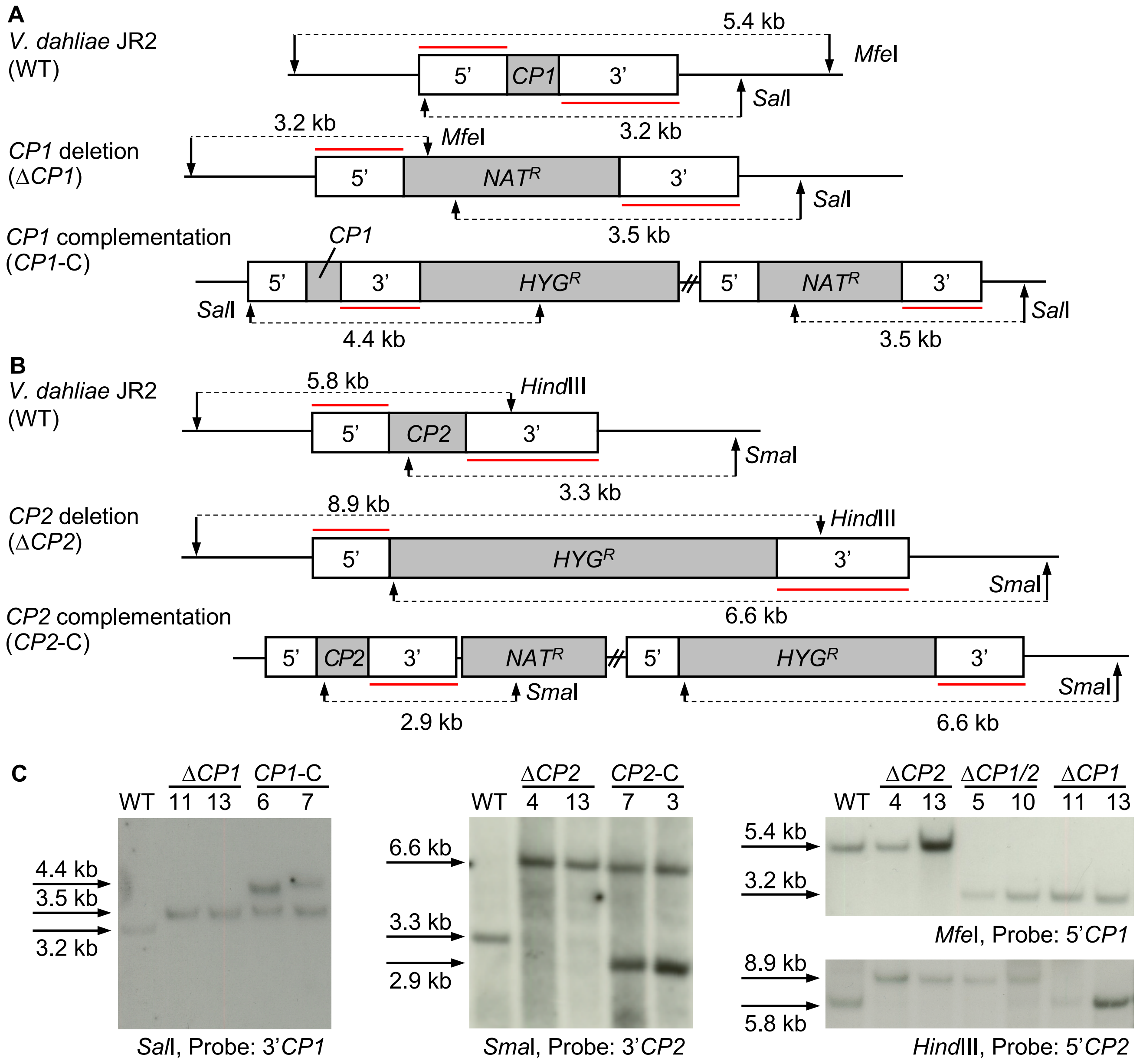
A
V. dahliae JR2 (WT)
5.4 kb
MfeI
5'
CP1
3'
SalI
3.2 kb
CP1 deletion (ΔCP1)
3.2 kb
MfeI
5'
NATR
3'
SalI
3.5 kb
CP1 complementation (CP1-C)
CP1
5'
3'
HYGR
5'
NATR
3'
SalI
SalI
4.4 kb
3.5 kb
B
V. dahliae JR2 (WT)
5.8 kb
HindIII
5'
CP2
3'
SmaI
3.3 kb
CP2 deletion (ΔCP2)
8.9 kb
HindIII
5'
HYGR
3'
SmaI
6.6 kb
CP2 complementation (CP2-C)
5'
CP2
3'
NATR
5'
HYGR
3'
SmaI
SmaI
2.9 kb
6.6 kb
C
WT
ΔCP1
11
13
CP1-C
6
7
4.4 kb
3.5 kb
3.2 kb
SalI, Probe: 3'CP1
WT
ΔCP2
4
13
CP2-C
7
3
6.6 kb
3.3 kb
2.9 kb
SmaI, Probe: 3'CP2
WT
ΔCP2
4
13
ΔCP1/2
5
10
ΔCP1
11
13
5.4 kb
3.2 kb
MfeI, Probe: 5'CP1
8.9 kb
5.8 kb
HindIII, Probe: 5'CP2

**S2 Figure**

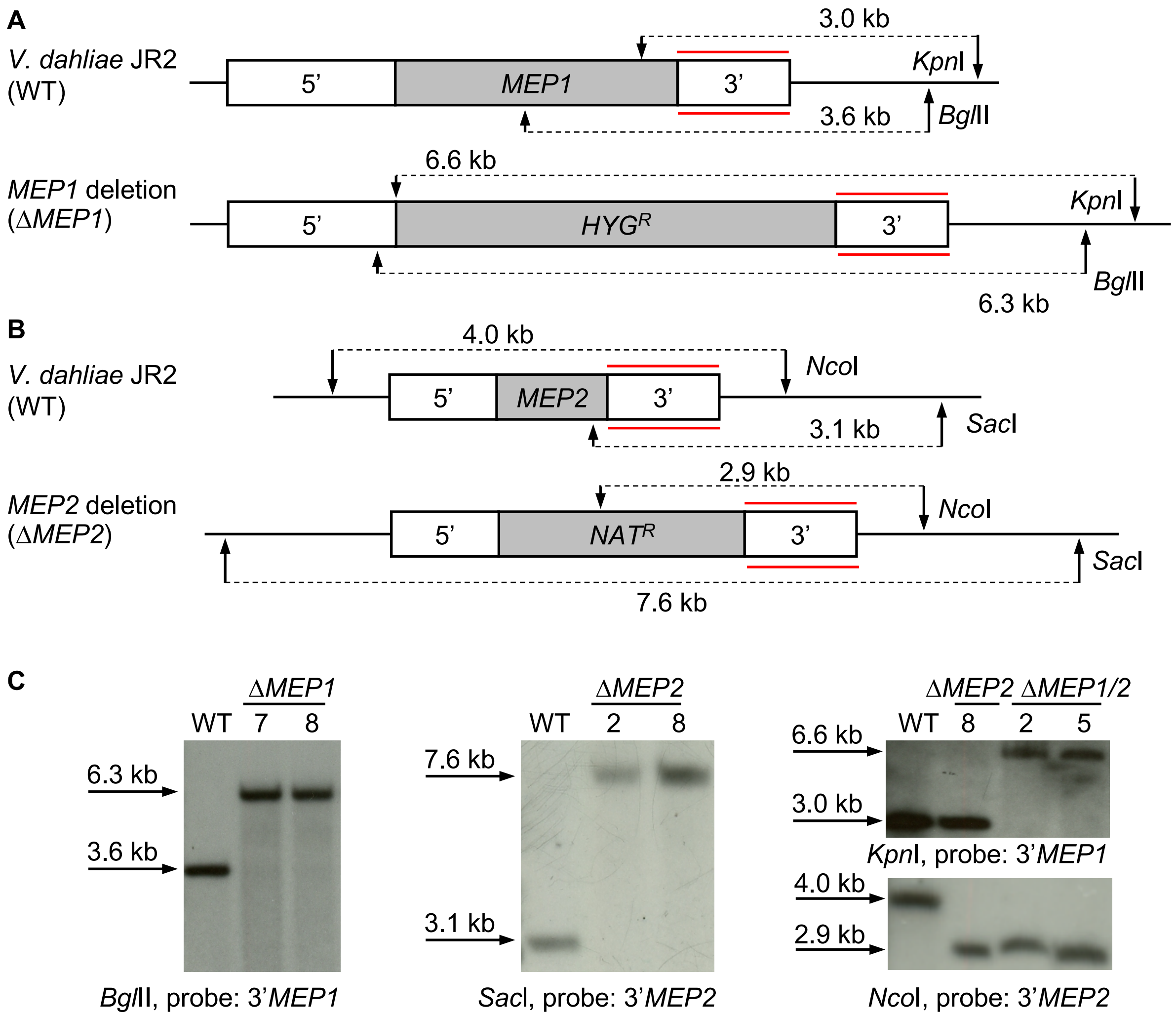
A
V. dahliae JR2 (WT)
3.0 kb
5'
MEP1
3'
KpnI
3.6 kb
BglII
MEP1 deletion (ΔMEP1)
6.6 kb
5'
HYGR
3'
KpnI
BglII
6.3 kb
B
V. dahliae JR2 (WT)
4.0 kb
NcoI
5'
MEP2
3'
3.1 kb
SacI
MEP2 deletion (ΔMEP2)
2.9 kb
5'
NATR
3'
NcoI
SacI
7.6 kb
C
ΔMEP1
WT 7 8
6.3 kb
3.6 kb
BglII, probe: 3'MEP1
ΔMEP2
WT 2 8
7.6 kb
3.1 kb
SacI, probe: 3'MEP2
ΔMEP2 ΔMEP1/2
WT 8 2 5
6.6 kb
3.0 kb
KpnI, probe: 3'MEP1
4.0 kb
2.9 kb
NcoI, probe: 3'MEP2

**S3 Figure**

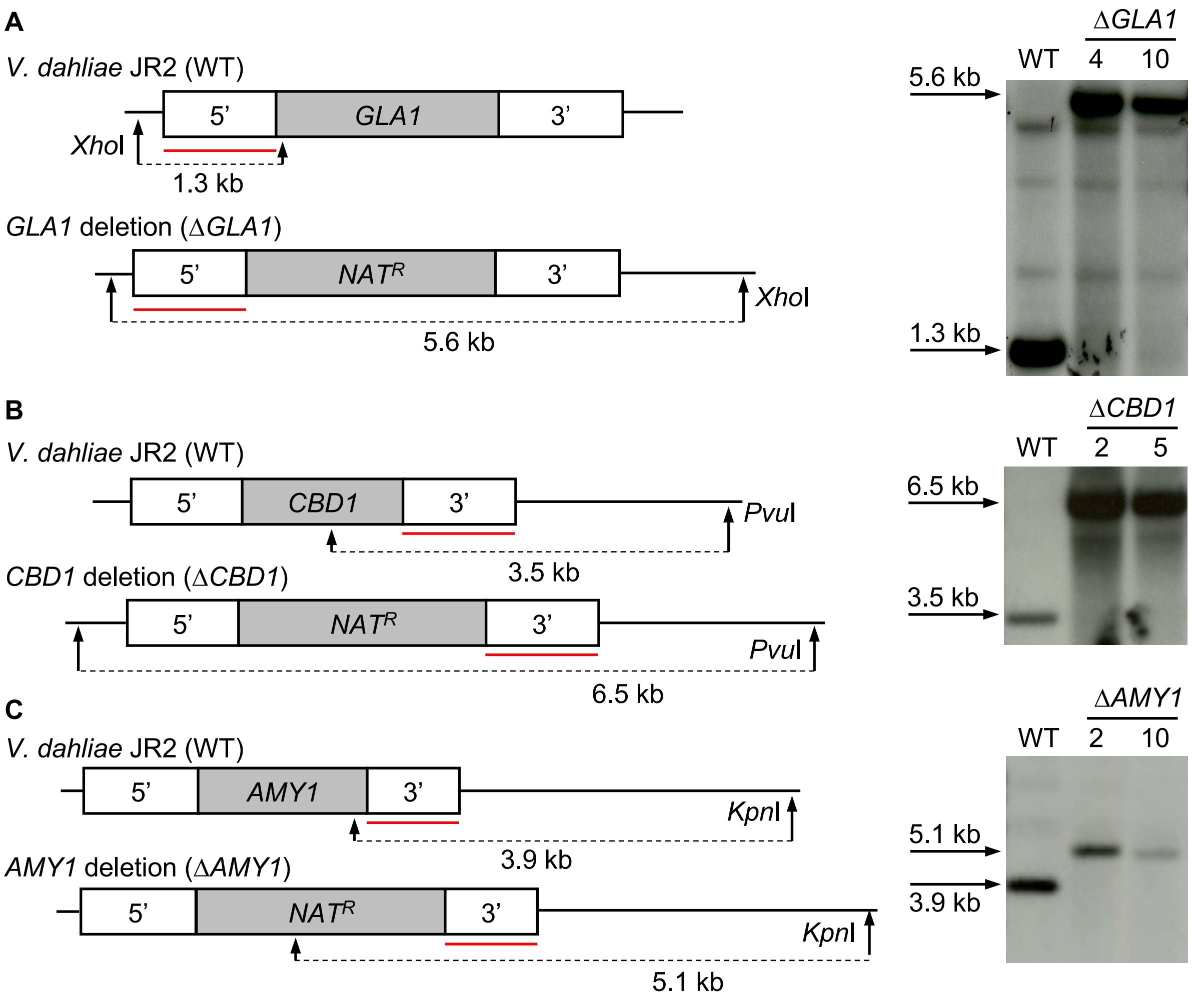
A
V. dahliae JR2 (WT)
5'
GLA1
3'
XhoI
1.3 kb
GLA1 deletion (ΔGLA1)
5'
NATR
3'
XhoI
5.6 kb
ΔGLA1
WT
4
10
5.6 kb
1.3 kb
B
V. dahliae JR2 (WT)
5'
CBD1
3'
PvuI
3.5 kb
CBD1 deletion (ΔCBD1)
5'
NATR
3'
PvuI
6.5 kb
ΔCBD1
WT
2
5
6.5 kb
3.5 kb
C
V. dahliae JR2 (WT)
5'
AMY1
3'
KpnI
3.9 kb
AMY1 deletion (ΔAMY1)
5'
NATR
3'
KpnI
5.1 kb
ΔAMY1
WT
2
10
5.1 kb
3.9 kb

**S4 Figure**

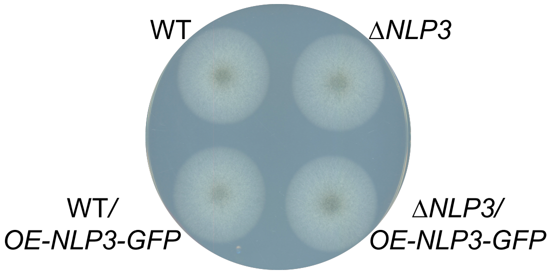


**S5 Figure**

**S1 Text. Additional Materials and Methods.**

**Gene deletion and overexpression in the haploid *V. dahliae***

For deletion of *NLP3* (*VDAG_JR2_Chr4g05950a*) ORF the 5' and 3' flanking regions were amplified from *V. dahliae* WT gDNA with ML86/ML87 (1 561 bp) and ML88/ML89 (1 284 bp), respectively. The nourseothricin resistance cassette was obtained by amplification of pME4815 with ML8/ML9 (2 194 bp). To receive the backbone, pME4564 was cut with *Stu*I and *Eco*RV and the linearized backbone (6 804 bp) was ligated with the fragments resulting in pME4883. *V. dahliae* JR2 WT was transformed with this construct resulting in *NLP3* deletion strains (VGB384, VGB385). Transformants were tested for correct integration of the construct by Southern hybridization (S1 Fig). The gDNA of WT and transformant strains was treated with *Xho*I and the 5' flanking region of *NLP3* was prepared as probe to verify the *NLP3* deletion strains.

Additionally, strains expressing free *GFP* or *NLP3-GFP* were constructed. The *NLP3* deletion strain VGB384 was transformed with pGreen2 (Tran *et* al., 2014) to generate the *NLP3* deletion strain expressing free *GFP*. Transformants were screened for their fluorescence signal by confocal microscopy after growing fungal hyphae in µ-slide 8 well microcopy chambers (Ibidi) at 25°C overnight. Selected transformants were next tested by Southern hybridization to possess the ectopically integrated fragments (S1 Fig). Genomic DNA of WT and the transformants was cut by *Bgl*I. Correct strains were verified with *GFP* as a probe. VGB431 and VGB432 were conserved. To overexpress C-terminally tagged *NLP3* in the WT or *NLP3* deletion strain, an *NLP3-GFP* possessing construct was generated including the hygromycin resistance cassette and integrated into WT or the *NLP3* deletion strain VGB384, respectively. The *gpdA* promoter was amplified from pGreen2 with ML99/RH635 (874 bp), the *NLP3* sequence without the stop codon was amplified from genomic WT DNA with ML100/ML101 (874 bp). ML101 includes a sequence coding for GGSGG that serves as a flexible linker to allow independent protein folding (van Rosmalen et al., 2017). These fragments were fused in another PCR using primer pair ML101/RH635. *GFP* (without the start codon) fused to *trpC* terminator was obtained by PCR of pGreen2 with RH686/RH636 (1 507 bp). These fragments were ligated to *Eco*RV linearized pPK2. The constructed plasmid was named pME4990. This plasmid was used to transform the desired fragment into *V. dahliae* WT and VGB384. Transformants were tested by Southern hybridization as VGB431 an VGB432 (S1 Fig). VGB407 and VGB408 were verified to possess the *NLP3-GFP* in the WT and VGB409 and VGB410 showed positive signals for the construct in the *NLP3* deletion strain.

To generate an *NLP2* (*VDAG_JR2_Chr2g05460a*) deletion strain, the 5' and 3' flanking regions were amplified with ML90/ML91 (975 bp) and ML92/ML106 (1 165 bp), respectively, from WT gDNA and the $HYG^R$ cassette with ML8/RO3 (3 942 bp) of pPK2 [1]. Plasmid pME4564 was cut with *Stu*I and *Eco*RV and the linearized backbone (6 804 bp) was ligated with the fragments resulting in pME4885. *V. dahliae* WT and *NLP3* deletion strain were transformed with the construct and by homologous recombination the *NLP2* deletion strain and *NLP2/NLP3* double deletions strains were generated, respectively. Transformants were verified by Southern hybridization (S1 Fig). Verification of *NLP2* single and *NLP2/3* double deletion strains was obtained by processing gDNA with *Xho*I and 5' *NLP3* and 5' *NLP2* were used as probes to confirm VGB390, VGB391, VGB400 and VGB401.

For construction of the *CP1* (*VDAG_JR2_Chr7g00860a*) deletion strain, the 5' and 3' flanking regions were amplified of WT gDNA with AO84/AO85 (891 bp) and AO86/AO87 (1 205 bp), respectively. The nourseothricin resistance cassette was fused together with 5' and 3'flanking regions in a fusion PCR with AO84/AO87 (4 291 bp), which was then phosphorylated with T4 Polynucleotide Kinase (Thermo Fisher Scientific). The backbone of pME4564 (6 804 bp) was received by enzyme restriction with *Eco*RV and *Stu*I. It was dephosphorylated with FastAP Thermosensitive Alkaline Phosphatase (Thermo Fisher Scientific) and ligated to the insert with T4 DNA Ligase. *V. dahliae* JR2 was transformed with the resulting plasmid pME4887 to generate the *CP1* deletion strain. For the construct of the complementation strain the *CP1* ORF was amplified including the 5' and 3' flanking regions with AO131/AO132 (2 626 bp) and ligated to *Eco*RV linearized pPK2 (10 751 bp) resulting in pME4888. The complementation was ectopically integrated into *CP1* deletion strain. To verify the constructed strains they were subjected to Southern hybridization (S2 Fig). For confirmation of *CP1* deletion strains VGB316 and VGB317 and *CP1* complementation strains VGB489 and VGB490 gDNA of WT and the transformants was cut with *Sal*I while 3' *CP1* region was used as probe.

Construction of *CP2* (*VDAG_JR2_Chr2g07000a*) deletion strain required amplification of 5' and 3' flanking region of *CP2* with ML94/ML95 (778 bp) and ML96/ML97 (1 343 bp), respectively, of WT gDNA. Together with the $HYG^R$ cassette the fragments were ligated to the backbone of pME4564, which was linearized with *Stu*I and *Eco*RV (6 804 bp). The resulting plasmid pME4889 was used to transform *V. dahliae* WT resulting in the *CP2* deletion strains VGB406 and VGB422. For the complementation the plasmid pME4890 was constructed by amplifying the *CP2* ORF including its 5' and 3' flanking regions with ML104/ML105 (2 913 bp) of WT gDNA, which was ligated to *Eco*RV linearized pME4815. This construct was utilized to transform *CP2* deletion strain VGB406 resulting in the ectopic complementation strains VGB429 and VGB430. Additionally, VGB406 was transformed with pME4887 to generate the Δ*CP1*Δ*CP2* double deletion strains VGB423 and VGB424. All strains were confirmed in Southern hybridizations (S2 Fig). To test *CP2* strains, gDNA was cut with *Sma*I and correct fragments were detected with 3' *CP2* region as probe resulting in the verification of *CP2* deletion and ectopic complementation strains. For verification of Δ*CP1*Δ*CP2* strains gDNA was cut with *Mfe*I or *Hind*III and as probes the 5' flanking region of *CP1* or *CP2* were utilized, respectively.

For the deletion of *MEP1* (*VDAG_JR2_Chr8g09760a*) the annotation of VdLs.17 *MEP1* (*VDAG_03418*) starting 341 bp upstream of the JR2 prediction was considered in addition to the prediction for *V. dahliae* JR2 to ensure the whole gene is deleted. The corresponding 5' and 3' flanking regions were amplified of WT gDNA with JST27/JST28 (1 500 bp) and JST29/JST30 (1 000 bp), respectively. The fragments were ligated together with the $HYG^R$ cassette to the pME4564 backbone, which was obtained by PCR amplification with ML1/ML2 (6 727 bp), resulting in pME4891. *V. dahliae* WT was transformed with the deletion construct and *MEP1* deletion strains VGB226 and VGB227 were confirmed by Southern hybridization (S3 Fig). Genomic DNA was cut with *Bgl*II and *MEP1* 3' flanking region was used as probe.

To generate the *MEP2* (*VDAG_JR2_Chr1g21900a*) deletion strain the 5' and 3' flanking regions were amplified with ML12/ML13 (960 bp) and ML14/ML15 (1 000 bp) of WT gDNA, respectively. Together with the $NAT^R$ cassette the fragments were ligated to the backbone of pME4564, which was generated by PCR amplification with ML1/ML2 (6 727 bp). The resulting plasmid pME4892 was used for construction of *MEP2* deletion strain by transformation of *V. dahliae* WT. Genomic DNA of transformants and WT was processed with *Sac*I and *MEP2* deletion strains VGB126 and VGB133 were confirmed by Southern hybridization with 3' *MEP2* as probes (S3 Fig). Strain VGB126 was further transformed with pME4891 to obtain the double deletion strains Δ*MEP1*Δ*MEP2* VGB203 and VGB204 that were verified by Southern hybridization with *Kpn*I treated gDNA and 3' *MEP1* or *Nco*I treated gDNA and 3' *MEP2*.

To obtain plasmids for the generation of deletion strains with a $NAT^R$ cassette, the 5' and 3' flanking regions of the respective genes were amplified of *V. dahliae* WT gDNA. Fragments were then ligated to the backbone of pME4564 that was gained by PCR amplification with ML1/ML2 (6 727 bp).

The *GLA1* (*VDAG_JR2_Chr8g11020a*) deletion construct pME4894 contains the 5' and 3' flanking regions amplified with ML20/ML21 (990 bp) and ML22/ML23 (1 095 bp), respectively. After transformation of *V. dahliae* WT with this construct, the resulting transformants were tested for their correct insert by Southern hybridization with *Xho*I treatment of the gDNA and 5' flanking region as probe (S4 Fig). The confirmed *GLA1* deletion strains VGB129 and VGB130 were conserved.

For the *CBD1* (*VDAG_JR2_Chr4g04440a*) deletion construct the 5' and 3' flanking regions were amplified with ML16/ML17 (982 bp) and ML18/ML19 (996 bp), respectively. The resulting plasmid pME4895 was used to generate *CBD1* deletion strains. VGB127 and VGB132 were confirmed by Southern hybridization (S4 Fig). The gDNA was cut with *Pvu*I and 3' flanking region of *CBD1* was used as probe.

Construction of *AMY1* (*VDAG_JR2_Chr7g03330a*) deletion construct involves amplification of the 5' and 3' flanking regions with ML24/ML25 (1 013 bp) and ML26/ML27 (814 bp), respectively. The construct pME4896 was utilized for transformation of *V. dahliae* WT resulting in *AMY1* deletion strains VGB128 and VGB130. The strains were verified by Southern hybridization with *Kpn*I treated gDNA and 3' flanking region as probe (S4 Fig).
